# *Plasmodium* sporozoites require the protein B9 to invade hepatocytes

**DOI:** 10.1101/2021.10.25.465731

**Authors:** Priyanka Fernandes, Manon Loubens, Carine Marinach, Romain Coppée, Morgane Grand, Thanh-Phuc Andre, Soumia Hamada, Anne-Claire Langlois, Sylvie Briquet, Philippe Bun, Olivier Silvie

## Abstract

*Plasmodium* sporozoites are transmitted to a mammalian host during blood feeding by an infected mosquito and invade hepatocytes for initial replication of the parasite in the liver. This leads to the release of thousands of merozoites into the blood circulation and initiation of the pathogenic blood stages of malaria. Merozoite invasion of erythrocytes has been well characterized at the molecular and structural levels. In sharp contrast, the molecular mechanisms of sporozoite invasion of hepatocytes are poorly characterized. Here we report a new role during sporozoite entry for the B9 protein, a member of the 6-cysteine domain protein family. Using genetic tagging and gene deletion approaches in rodent malaria parasites, we show that B9 is secreted from sporozoite micronemes and is required for productive invasion of hepatocytes. Structural modelling indicates that the N-terminus of B9 forms a beta-propeller domain structurally related to CyRPA, a cysteine-rich protein forming an invasion complex with Rh5 and RIPR in *P. falciparum* merozoites. We provide evidence that the beta-propeller domain of B9 is essential for protein function during sporozoite entry and interacts with P36 and P52, both also essential for productive invasion of hepatocytes. Our results suggest that, despite using distinct sets of parasite and host entry factors, *Plasmodium* sporozoites and merozoites may share common structural modules to assemble protein complexes for invasion of host cells.

## INTRODUCTION

Malaria is caused by *Plasmodium* spp. parasites and still remains a major health and socio-economic problem in endemic countries^1^. Sporozoites, the mosquito-transmitted forms of the malaria parasite, first infect the liver for an initial and obligatory round of replication, before initiating the symptomatic blood stages. Infection of the liver is clinically silent and constitutes an ideal target for a malaria vaccine. Until now, only one single antigen, the circumsporozoite protein (CSP), had been considered for clinical vaccine development against the extracellular sporozoite stage, with limited success^2^. Other sporozoite antigens, especially parasite proteins involved in host-parasite interactions, could be considered as potential vaccine targets to prevent sporozoite entry into hepatocytes. This highlights the need to better characterize the molecular mechanisms of sporozoite infection in order to identify new vaccine targets.

Like other Apicomplexan parasites, *Plasmodium* invades host cells using a unique mechanism that involves the sequential secretion of apical organelles, called micronemes and rhoptries, and the formation of a moving junction (MJ) through which the parasite actively glides to enter the cell and form a specialized parasitophorous vacuole (PV) where it further replicates^3^. Proteins released from micronemes onto the parasite surface are prime candidates to interact with host cell surface receptors, triggering subsequent secretion of the rhoptry content, formation of the MJ and commitment to productive invasion. However, until now the ligand-receptor interactions mediating *Plasmodium* sporozoite invasion and the nature of the sporozoite MJ have remained enigmatic^4^.

We previously characterized host entry pathways used by human (*P. falciparum*, *P. vivax*) and rodent (*P. yoelii*, *P. berghei*) parasites to infect hepatocytes^5, 6^, and showed that CD81 and SR-BI define independent entry routes for *P. falciparum* and *P. vivax* sporozoites, respectively^6^. Remarkably, this alternative usage of host cell receptors is also observed with rodent malaria model parasites, providing robust and tractable experimental systems^6, 7^. Indeed, *P. yoelii* sporozoites, like *P. falciparum*, strictly require CD81 to infect liver cells, whereas *P. berghei* can alternatively use CD81 or SR-BI for productive invasion^6^. Only two parasite proteins, P36 and P52, have been identified as being specifically required for productive invasion of hepatocytes^6, 8–11^. Using inter-species genetic complementation in mutant *P. berghei* and *P. yoelii* lines, we showed that P36 is a key determinant of host cell receptor usage, establishing for the first time a functional link between sporozoite and host cell entry factors^6^. The molecular function of P36 remains unknown. One study proposed that P36 interacts with the ephrin receptor EphA2 on hepatocytes to mediate infection^12^, but direct evidence for such an interaction is lacking, and EphA2 was later shown to be dispensable for sporozoite productive invasion^13^. Interestingly, interspecies genetic complementation experiments showed that *P. berghei* Δ*p52*Δ*p36* mutants complemented with PyP52 and PyP36 exhibit a *P. yoelii*-like phenotype as they preferentially infect CD81-expressing cells^6^. However, whilst *P. yoelii* sporozoites are unable to infect hepatocytes in the absence of CD81, complemented *P. berghei* mutants retain a residual invasion capacity in CD81-deficient cells^6^. Furthermore, genetic complementation with *P. falciparum* or *P. vivax* P52 and P36 cannot restore infection of Δ*p52*Δ*p36 P. berghei* sporozoites^6^. These results strongly suggest that additional parasite factors contribute to receptor-dependent productive invasion.

P36 and P52 both belong to the so-called 6-cysteine domain protein family, which is characterized by the presence of one or several 6-cysteine (6-cys) domains^14^. 6-cys domains are ∼120 amino acid long domains containing four or six conserved cysteine residues that respectively form two or three disulphide bonds resulting in a beta-sandwich fold^14^. *Plasmodium* spp. possess 14 members of the 6-cys protein family^15^. *Plasmodium* 6-cys proteins are typically expressed in a stage-specific manner, and have been implicated in protein-protein interactions in *P. falciparum* merozoites^16, 17^, gametocytes^18, 19^, ookinetes^20^ and sporozoites^11^. Proteomic studies have shown that, in addition to P36 and P52, *Plasmodium* sporozoites express three other 6-cys proteins, P12p, P38 and B9^21–24^. While the contribution of P12p and P38 had not been studied until now, a previous study reported that the protein B9 is not expressed in sporozoites due to translational repression, and is not required for sporozoite invasion but is needed during infection of hepatocytes for early maintenance of the PV^15^.

Here, we systematically analysed the role of P12p, P38 and B9 during sporozoite invasion, using a reverse genetics approach based on our Gene Out Marker Out (GOMO) strategy^25^. We report that *b9* gene deletion totally abrogates sporozoite infectivity, whilst *p12p* and *p38* are dispensable for hepatocyte infection in both *P. berghei* and *P. yoelii*. We show that B9 is a sporozoite micronemal protein and that B9-deficient sporozoites fail to productively invade hepatocytes. Secondary structure analysis and protein structure modelling indicate that B9 is a hybrid protein containing a CyRPA-like beta propeller domain in addition to non-canonical 6-cys domains. Structure-guided mutagenesis reveals that the propeller domain is not associated with host receptor usage but is essential for B9 function, possibly through the assembly of supramolecular protein complexes with the 6-Cys proteins P36 and P52 during host cell invasion.

## RESULTS

### Analysis of the repertoire of *Plasmodium* sporozoite 6-cys proteins suggests that P36, P52 and B9 are employed by infectious sporozoites only

In order to define the repertoire of 6-cys proteins expressed at the sporozoite stage, we first analysed the proteome datasets of *P. falciparum*^22, 26^, *P. vivax*^24^, *P. yoelii*^26^ and *P. berghei*^21^ sporozoites. As expected, P36 and P52 were identified by mass spectrometry in sporozoites from all four species. Interestingly, three other 6-cys proteins, P12p, P38 and B9, were consistently identified across the datasets. Among this core of five 6-cys proteins, P12p and P38 have been identified in the surface proteome of *P. falciparum* sporozoites, P12p being quantitatively enriched on the surface of activated parasites in the presence of bovine serum albumin^27^. Interestingly, P12p and P38 do not seem to be uniquely employed by sporozoites, as they have been detected in *P. falciparum* asexual and sexual blood stages^28–32^, and in *P. berghei* gametocytes^33^, respectively. In contrast, P36, P52 and B9 were identified in sporozoites only, and a recent study identified P36, P52 and B9 as up-regulated in infectious sporozoites (UIS) proteins in *P. falciparum* and *P. yoelii*, whilst P12p and P38 were also detected in oocyst-derived sporozoites^34^. These observations suggest that B9, like P36 and P52, may play a role in mature sporozoites.

### Reverse genetics analysis in rodent malaria parasites shows that *b9* (but not *p12p* and *p38*) is essential for sporozoite infectivity

A previous study reported that B9 is not expressed in sporozoites and is required for early liver stage development but not host cell invasion^15^. The contribution of P12p and P38 during sporozoite invasion has not been investigated so far, although the *p38* gene could be deleted in *P. berghei* without any detectable phenotypic defect during blood stage parasite growth and transmission to mosquitoes^35, 36^. Given the consistent detection of P12p, P38 and B9 proteins in sporozoites by mass spectrometry, we sought to determine the functional importance of these proteins in *P. berghei* and *P. yoelii* sporozoites using a reverse genetics approach. We used our GOMO strategy^37^ to replace genes of interest, through homologous recombination with a GFP expression cassette under the control of a constitutive HSP70 promoter, to facilitate monitoring of host cell invasion (**Supplementary Fig. 1a**). Targeting vectors were assembled by inserting 5’ and 3’ homology fragments of *P. berghei* or *P. yoelii p12p* (PBANKA_0111100; PY17X_0112700), *p38* (PBANKA_1107600; PY17X_1108700) and *b9* (PBANKA_0808100; PY17X_0811300) genes in the GOMO-GFP plasmid^37^, and used to transfect wild type (WT) *P. berghei* (ANKA) or *P. yoelii* (17XNL) blood stage parasites. We then applied the GOMO selection strategy, consisting of positive selection with pyrimethamine, negative selection with 5-fluorocytosine and flow cytometry-assisted parasite sorting, as previously described^37^. Pure populations of GFP-expressing drug-selectable marker-free PbΔ*p12p*, PbΔ*p38*, PbΔ*b9*, PyΔ*p12p*, PyΔ*p38* and PyΔ*b9* parasite lines were obtained, confirming that none of the targeted genes are essential during blood stage replication of the parasite. Genotyping by PCR confirmed gene deletion and excision of the drug-selectable marker cassette, as desired, in all parasite lines (**Supplementary Fig. 1b-h**). All the mutants could be transmitted to mosquitoes and produced normal numbers of salivary gland sporozoites, similar to Δ*p36* parasites (**Fig. 1a-b**). We then assessed the infectivity of the *P. berghei* and *P. yoelii* mutant lines in C57BL/6 and BALB/c mice, respectively. C57BL/6 mice injected with 10,000 PbΔ*p12p* or PbΔ*p38* sporozoites all developed a patent blood stage infection, like the parental PbGFP parasites (**Fig. 1c**). Similarly, BALB/c mice injected with 10,000 PyΔ*p12p* or PyΔ*p38* sporozoites all developed a patent blood stage infection (**Fig. 1d**). In sharp contrast, none of the animals injected with *P. berghei* or *P. yoelii* Δ*b9* sporozoites developed parasitemia, phenocopying the Δ*p36* mutants (**Fig. 1c-d**). Abrogation of Δ*b9* sporozoite infectivity was also observed *in vitro* in hepatocyte cell lines. FACS analysis 24 hours post-infection revealed a dramatic reduction in the number of PbΔ*b9* exoerythrocytic forms (EEFs) in comparison to control PbGFP or PbΔ*p12p* and PbΔ*p38* sporozoites in HepG2 cells, which was similar to the reduction observed with PbΔ*p36* mutants (**Fig. 1e**). Using antibodies specific for UIS4, a marker of the PV membrane (PVM) that specifically labels productive vacuoles^38, 39^, we confirmed that, in contrast to Δ*p12p* and Δ*p38* mutants, Δ*b9* parasites were not able to form productive vacuoles (**Fig. 1f**). Together, these results show that *b9* is essential for sporozoite liver infection both *in vivo* and *in vitro*, corroborating the results of a previous study^15^, and that *p12p* and *p38* genes on the contrary are dispensable for parasite invasion and liver stage development.

**Figure 1.**
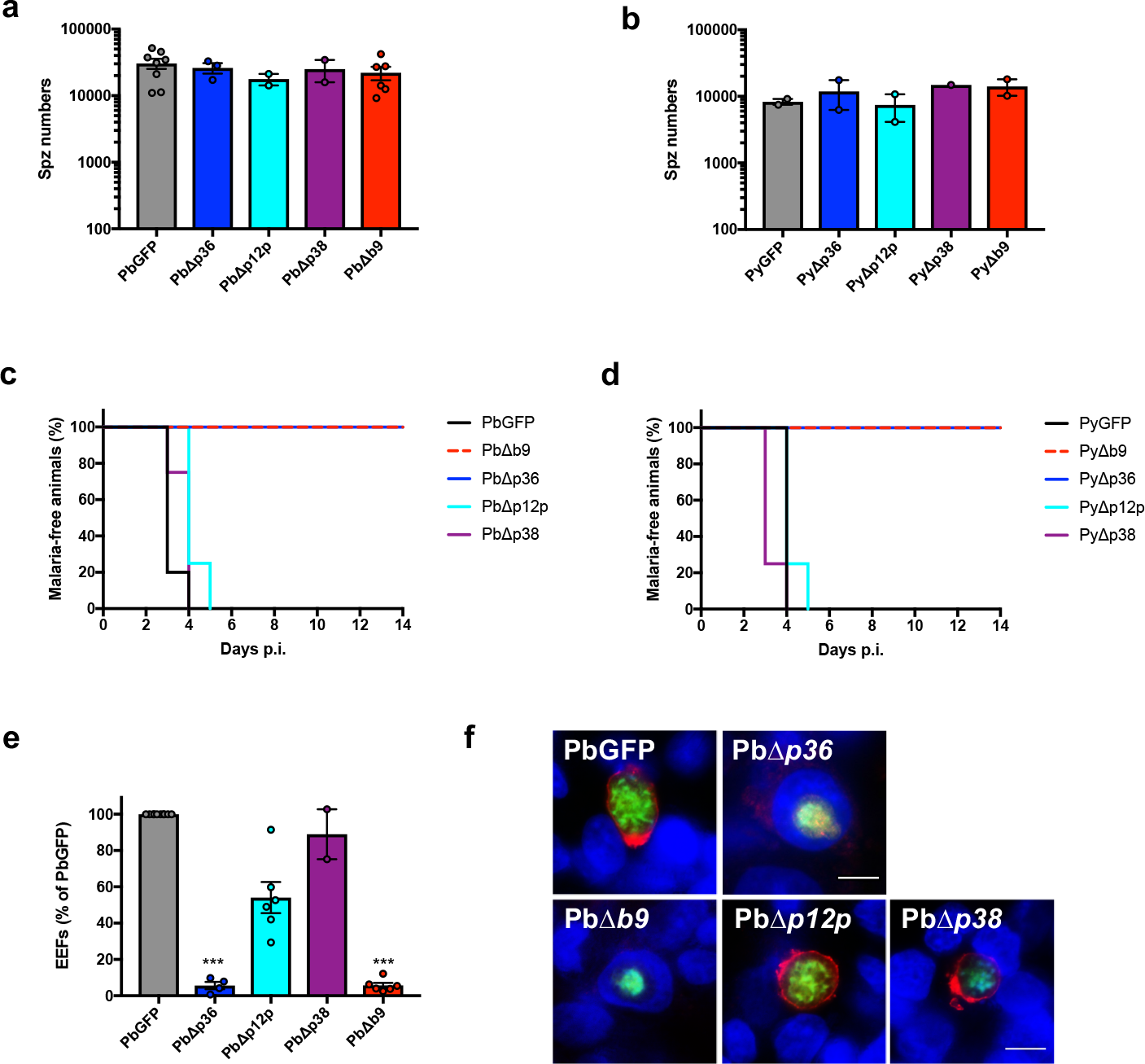
Deletion of *b9* but not *p12P* or *p38* genes abrogates sporozoite infectivity in *P. berghei* and *P. yoelii*. **a**, Number of sporozoites isolated from the salivary glands of mosquitoes infected with PbGFP, PbΔ*p36*, PbΔ*p12p,* PbΔ*p38* or PbΔ*b9* parasites (mean +/- SD; p = 0.67, one-way ANOVA). **b,** Number of sporozoites isolated from the salivary glands of mosquitoes infected with PyGFP, PyΔ*p36*, PyΔ*p12p*, PyΔ*p38* or PyΔ*b9* parasites (mean +/- SD; p = 0.66, one-way ANOVA). **c**, Kaplan-Meier analysis of time to patency in C57BL/6 mice (n = 5) after intravenous injection of 10^4^ PbGFP, PbΔ*p36*, PbΔ*p12p,* PbΔ*p38* or PbΔ*b9* sporozoites. Mice were followed daily for the appearance of blood stage parasites (p = 0.0001, Log rank Mantel-Cox test). **d**, Kaplan-Meier analysis of time to patency in BALB/c mice (n = 5) after intravenous injection of 10^4^ PyGFP, PyΔ*p36*, PyΔ*p12p*, PyΔ*p38* or PyΔ*b9* sporozoites. Mice were followed daily for the appearance of blood stage parasites (p < 0.0001, Log rank Mantel-Cox test). **e**, Infection rates were determined by quantification of EEFs (GFP-positive cells) 24 h after infection of HepG2 cell cultures with PbGFP, PbΔ*p36*, PbΔ*p12p,* PbΔ*p38* or PbΔ*b9* sporozoites. Results are expressed as % of control (PbGFP). ***p < 0.001 as compared to PbGFP (one-way ANOVA followed by Dunnett’s multiple comparisons test). **f**, Immunofluorescence images of HepG2 cells infected with PbGFP, PbΔ*p36*, PbΔ*p12p,* PbΔ*p38* or PbΔ*b9* parasites expressing GFP (green) and labelled with anti-UIS4 antibodies (red) and Hoechst 77742 (blue). PbGFP, PbΔ*p12p* and PbΔ*p38* are surrounded by a UIS4-positive PV membrane (red), while PbΔ*p36* and PbΔ*b9* parasites are intranuclear and lack a UIS4-positive PVM. Scale bar, 10 μm.

### B9 is required for sporozoite invasion

After infection of cell cultures with Δ*b9* sporozoites, only very low numbers of intracellular parasites were observed, all of which were seemingly intranuclear and lacked a UIS4-labeled PVM, similar to the Δ*p36* mutants (**Fig. 1f**). Intranuclear EEFs are known to result from cell traversal events^40^. Accordingly, a cell wound-repair assay confirmed that the cell traversal activity of Δ*b9* sporozoites is not different to PbGFP parasites, in both HepG2 and HepG2/CD81 cells (**Fig 2a-b**). In contrast, direct quantification of invaded cells by FACS revealed that host cell invasion by Δ*b9* sporozoites is greatly impaired in both cell types (**Fig. 2c-d**). Based on the similar phenotype observed with Δ*b9* and Δ*p36* parasites^6^, we hypothesized that B9 could play a role during productive invasion. Productive host cell invasion is associated with discharge of the sporozoite rhoptries, resulting in depletion of the rhoptry proteins RON2 and RON4^3, 41^. To monitor rhoptry discharge in B9-deficient parasites, we genetically modified the *ron4* locus in the PbΔ*b9* mutant line to replace the endogenous RON4 by a RON4-mCherry fusion by double homologous recombination (**Supplementary Fig. 2**). In parallel, we also genetically modified parental PbGFP and mutant PbΔ*p36* parasites, using the same RON4-targeting vector (**Supplementary Fig. 2**). Examination of PbGFP/*RON4-mCherry*, PbΔ*b9/RON4-mCherry* and PbΔ*p36/RON4-mCherry* by fluorescence microscopy confirmed expression of the rhoptry marker in merozoites and sporozoites, as expected^41^ (**Fig. 2e**). We then performed invasion assays in HepG2 cells and analysed the presence of the RON4-mCherry rhoptry marker by fluorescence microscopy. As expected, the RON4-mCherry signal was lost in a vast majority of intracellular PbGFP/*RON4-mCherry* sporozoites, reflecting rhoptry discharge during productive invasion (**Fig. 2f**). In sharp contrast, RON4-mCherry was detected in all examined PbΔ*b9* and PbΔ*p36* intracellular sporozoites, indicating that sporozoites lacking B9 or P36 invade cells without secreting their rhoptries, i.e. through traversal mode only. Altogether, these data demonstrate that genetic deletion of B9 abrogates productive host cell invasion by sporozoites, phenocopying the lack of P36. Our data also show that B9, like P36, is essential for both CD81-dependent and CD81-independent sporozoite entry.

**Figure 2.**
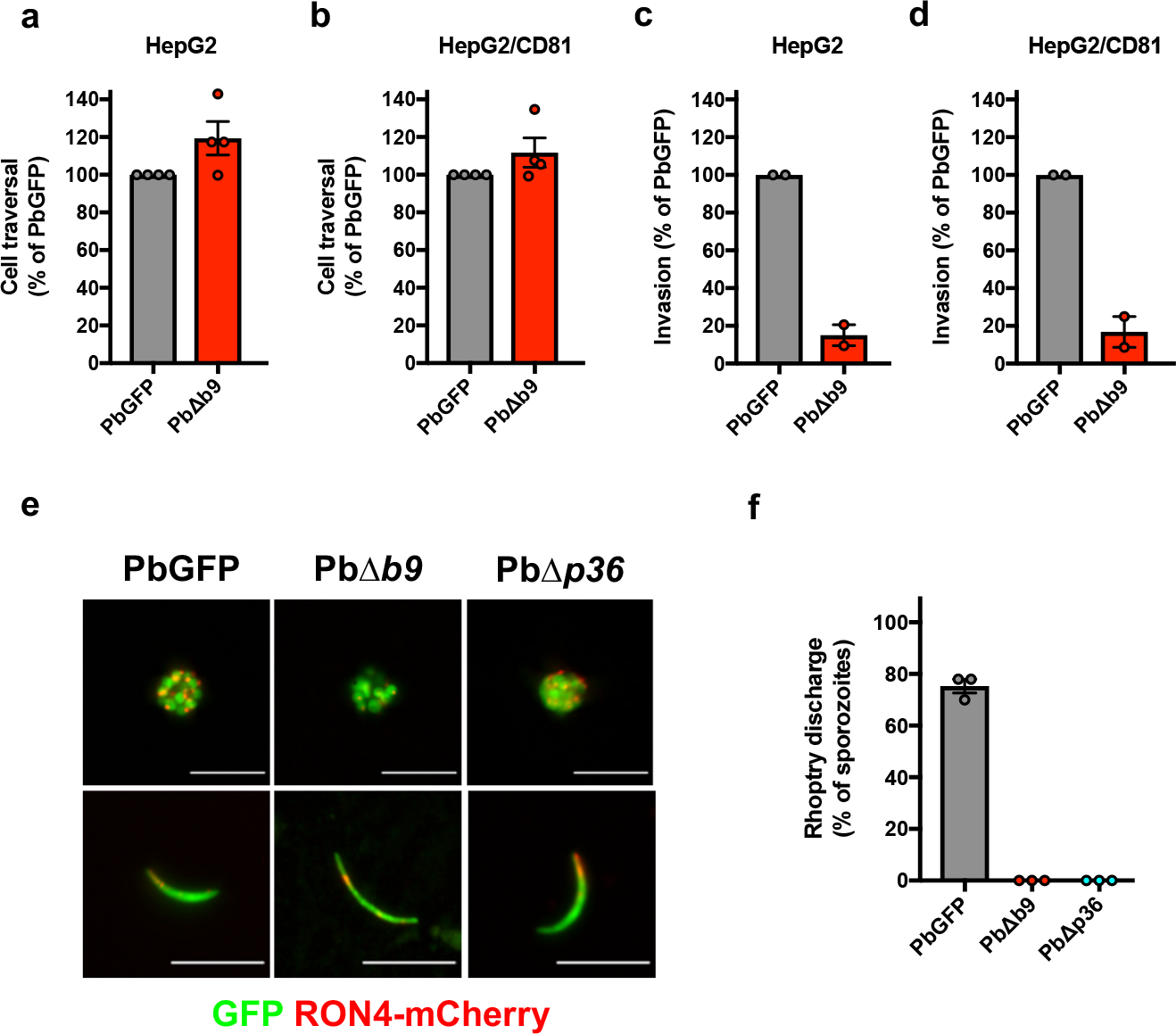
Sporozoites require B9 for productive invasion of host cells. **a-b**, Sporozoite cell traversal activity was analysed in HepG2 (a) and HepG2/CD81 (b) cell cultures incubated for 3 hours with PbGFP or PbΔ*b9* sporozoites in the presence of rhodamine-labelled dextran. The number of traversed (dextran-positive) cells was determined by FACS. **c-d,** Sporozoite invasion rates were determined in HepG2 (c) and HepG2/CD81 (d) cell cultures incubated for 3 hours with PbGFP or PbΔ*b9* sporozoites. The percentage of invaded (GFP-positive) cells was determined by FACS. **e**, Fluorescence microscopy images of RON4-mCherry-expressing PbGFP, PbΔ*b9* and PbΔ*p36* erythrocytic schizonts (upper panels) and salivary gland sporozoites (lower panels). Scale bar, 10 μm. **f**, Rhoptry discharge was analysed by fluorescence microscopy examination of HepG2 cells incubated for 3 hours with RON4-mCherry-expressing PbGFP, PbΔ*b9* or PbΔ*p36* sporozoites. Results are expressed as the percentage of parasites without detectable RON4-mCherry signal.

### B9 is secreted from the sporozoite micronemes

The phenotype of Δ*b9* mutants, combined with proteomic data, implies that the protein B9 is expressed in *P. berghei* sporozoites and plays a crucial role during host cell productive invasion, unlike previously thought^15^. In order to confirm the expression of B9 at the protein level and define its localization, we genetically modified the endogenous *b9* locus in *P. berghei* (PbGFP) to insert a triple Flag epitope in the protein-coding sequence, through double homologous recombination (**Supplementary Fig. 3a**). Because B9 is predicted to be glycosylphosphatidylinositol (GPI) anchored, we inserted the 3xFlag tag towards the C-terminus of the protein, downstream of the putative 6-cys domains but upstream of the predicted omega site (aspartate residue at position 826). Correct integration of the construct was confirmed by PCR on genomic DNA from B9-Flag blood stage parasites (**Supplementary Fig. 3b**). Importantly, we observed no defect in sporozoite development (**Fig. 3a**) and infectivity (**Fig. 3b**) in the B9-Flag line, demonstrating that the insertion of a 3xFlag epitope in B9 sequence had no detrimental effect on the protein function.

**Figure 3.**
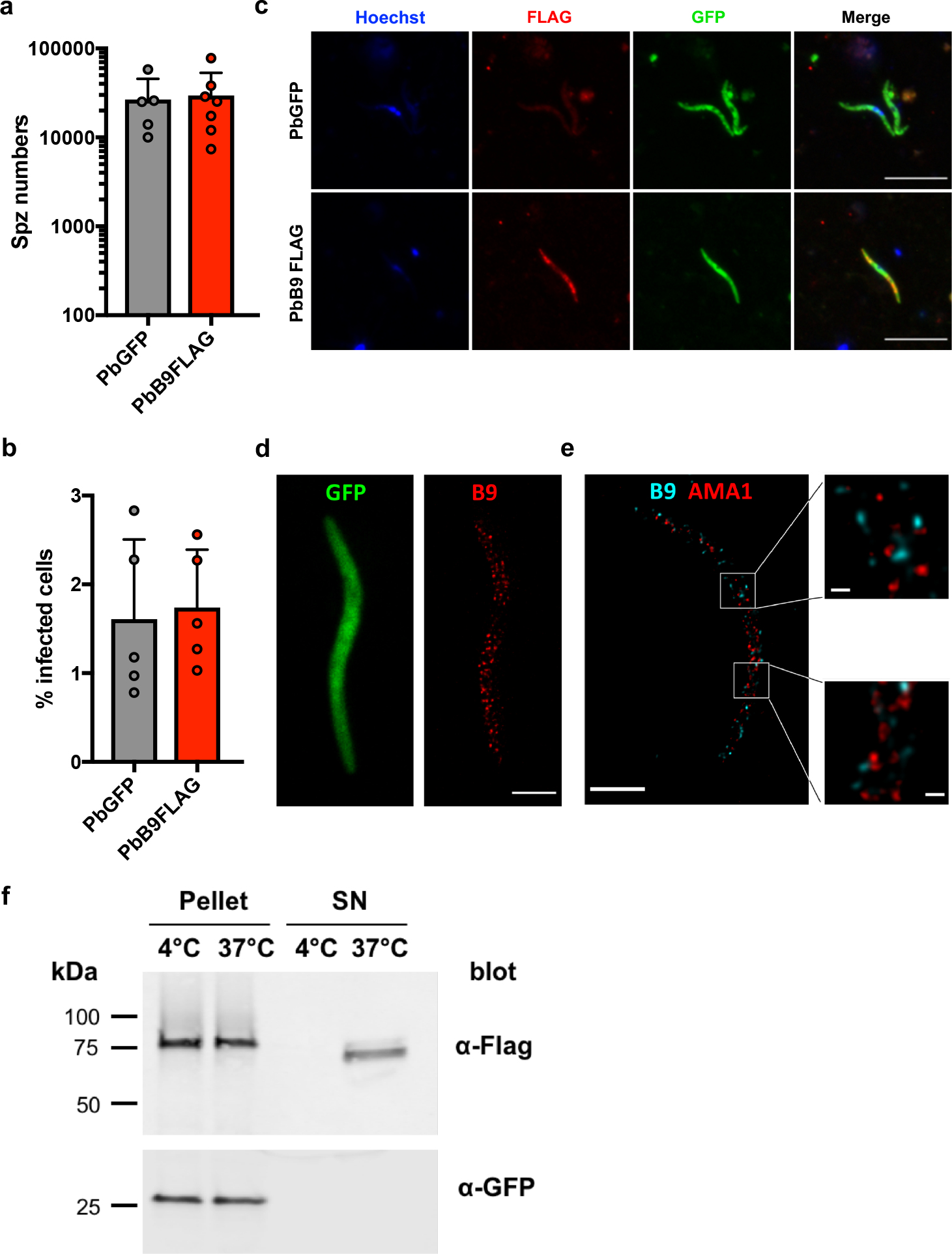
B9 localizes to a subset of sporozoite micronemes and is secreted upon parasite activation. **a**, Number of sporozoites isolated from the salivary glands of mosquitoes infected with PbGFP or PbB9-Flag parasites (mean +/- SD; p = 0.2893, Two-tailed ratio paired t test). **b,** Infection rates of PbGFP and PbB9-Flag parasites were determined in HepG2 cells 24 hours post-infection. The results show the percentage of invaded (GFP-positive) cells as determined by FACS (mean +/- SD; p = 0.6768, Two-tailed ratio paired t test). **c**, Immunofluorescence analysis of PbGFP and PbB9-Flag sporozoites labelled with anti-Flag antibodies (red). Parasites express GFP (green) and nuclei were stained with Hoechst 77742 (blue). Scale bar, 10 μm. **d**, Localization of B9 in sporozoites. First panel, confocal image of GFP (green); second panel, visualization of B9-Flag (red) using 2D STED (maximum intensity projection). Scale bar, 2 μm. **e**, STED images of a B9-flag sporozoite labelled with anti-AMA1 (red) and anti-Flag (cyan) antibodies. Scale bar, 2 μm (200 nm in insets). **f**, Immunoblot of B9-Flag sporozoite pellets and supernatants in control conditions (4°C) or after stimulation of microneme secretion (37°C), using anti-Flag or anti-GFP antibodies. The data shown are representative of three independent experiments.

Immunofluorescence with anti-Flag antibodies revealed that B9 is readily detected in B9-Flag salivary gland sporozoites, with a distribution pattern typical of a micronemal protein (**Fig. 3c**). Super-resolution microscopy using stimulated-emission-depletion (STED) showed that B9 distributes in numerous vesicles localized on each side of the nucleus, consistent with B9 being a micronemal protein (**Fig. 3d**). Interestingly, B9 did not co-localize with the apical membrane antigen 1 (AMA1), suggesting that the two proteins are present in distinct microneme subsets in salivary gland sporozoites (**Fig. 3e**). We next analysed the fate of B9 upon activation of sporozoite microneme secretion, by western blot. In non-activated control parasites, B9 was detected as a single band around 80 kDa (**Fig. 3f**). Upon stimulation of microneme secretion, B9 was also recovered in the supernatant fraction as a slightly smaller band, indicating that B9 is secreted from sporozoites upon activation, possibly after proteolytic processing (**Fig. 3f**). We failed to detect B9 on the surface of B9-Flag sporozoites by immunofluorescence, irrespective of parasite activation, suggesting that following microneme secretion, B9 is mainly released as a shed protein.

### B9 contains a CyRPA-like beta propeller domain

To get more insights into B9 properties, we investigated sequence and structural features of the protein using *P. falciparum* B9 as the reference sequence. Both hydrophobic cluster analysis and secondary structure prediction of B9 suggested that the whole sequence contains some strand and helix structures (**Supplementary Fig. 4**). However, no annotated conserved domain was detected at the sequence level using InterPro. In sharp contrast, three domains were predicted at the structural level using HHpred: an N-terminus propeller domain similar to that of CyRPA (e-value: 5.4e-03) encoded by the first exon, and two putative but poorly supported 6-cys domains encoded by the second exon (e-value > 1) (**Fig. 4a**). CyRPA is a cysteine-rich protein expressed in *P. falciparum* merozoites, where it forms a protein complex that is essential for invasion of erythrocytes^42, 43^. B9 is enriched in cysteines, nine being located in the predicted propeller domain that we suppose are involved in the formation of disulphide bonds in a similar manner to CyRPA^44^, to stabilize the protein structure (**Fig. 4a**).

**Figure 4.**
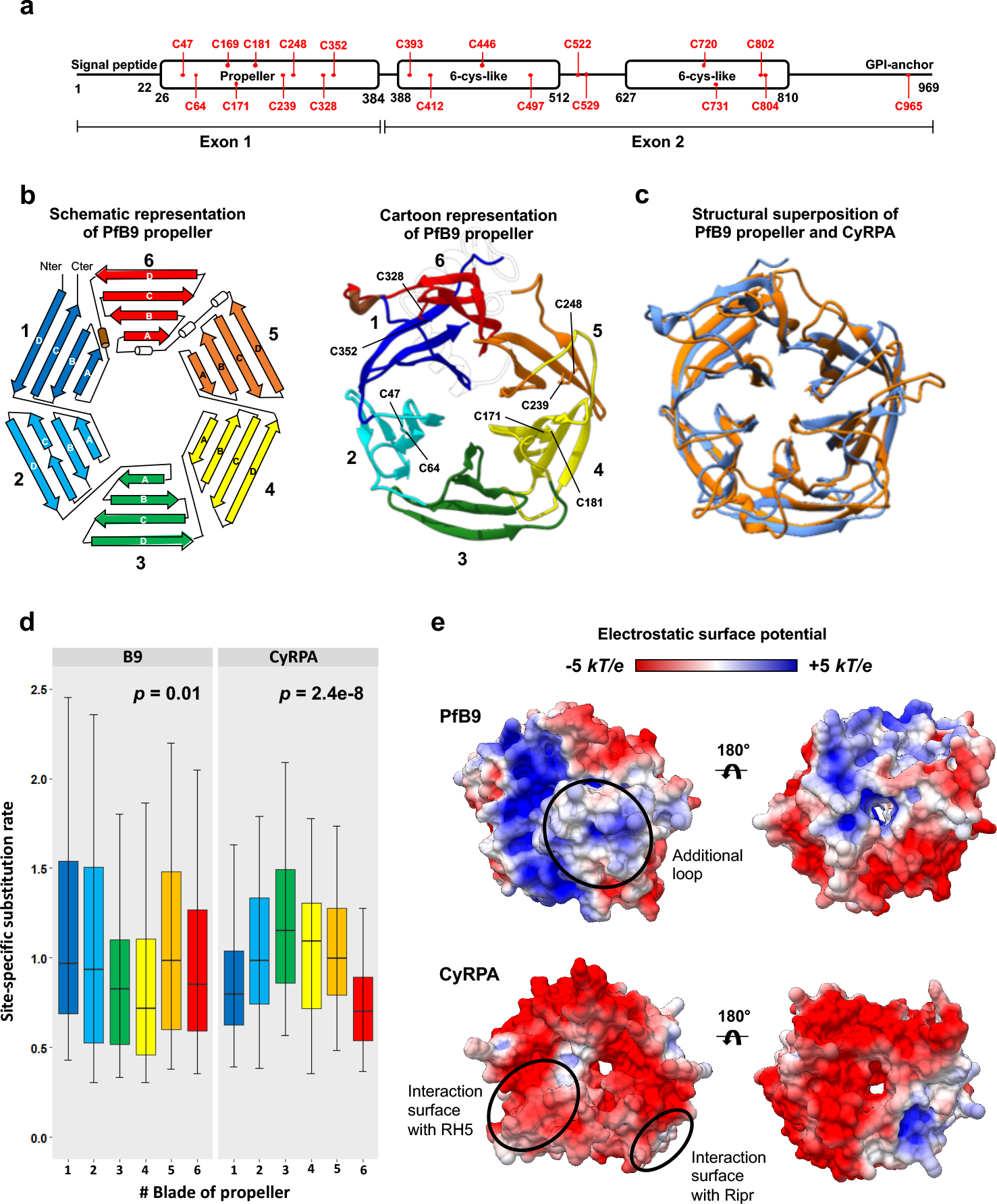
Structural and evolutionary features of B9 propeller. **a**, Predicted B9 conserved domains. PfB9 was used as the reference sequence. Cysteines are indicated in red. The delimitation of the domains is based on the HHpred results. B9 is composed of two exons, the first one covering the whole propeller domain. **b,** Predicted tertiary structure of PfB9 propeller. The predicted model is indicated as a schematic representation (*left*) and as cartoon (*right*). Each of the six blades is indicated with a specific color, labeled 1 to 6, and is composed of four-stranded anti-parallel beta-sheet, labeled A to D. The four disulfide bridges found in PfB9 are indicated. The long loop connecting blades 5 and 6 in the cartoon representation is transparent for ease of representation. **c,** Structural superposition of PfB9 propeller with CyRPA. PfB9 and CyRPA are respectively colored in blue and orange. Both superposition and RMSD calculation were based on all Cɑ atoms using the *MatchMaker* function in UCSF Chimera. **d,** Conservation level of the six blades of B9 propeller and CyRPA. Site-specific rates were estimated using the GP4Rate tool, and were compared between the six blades using non-parametric Kruskal-Wallis *H* test. Box boundaries represent the first and third quartiles and the length of whiskers correspond to 1.5 times the interquartile range. **e,** Electrostatic surface potential of PfB9 propeller and CyRPA structures, estimated with the APBS method. Electrostatic potential values are in units of kT/e at 298 K, on a scale of −5 kT/e (red) to +5 kT/e (blue). White color indicates a neutral potential. The missing charges were added using the Add Charge function implemented in USCF Chimera. The additional long loop connecting blades 5 and 6 of PfB9 propeller and the interaction surfaces of CyRPA with Rh5 and Ripr are indicated with circles.

To explore the structural features of the B9 propeller, we predicted the tertiary structure of PfB9 propeller (covering positions 26 to 386) by homology modelling using CyRPA as a template structure^44^ (PDB ID: 5TIH; **Supplementary File 1**; **Supplementary Fig. 5**). As expected, PfB9 adopted a six-bladed propeller structure, with each blade being composed of four-stranded anti-parallel beta-sheets (**Fig. 4b**). Four disulphide bonds were predicted within the blades which may stabilize each individual blade of the PfB9 propeller (C47-C64, C171-C181, C239-C248, C328-C352; **Fig. 4b**). Furthermore, a long loop connecting blades 5 and 6 and containing three putative short helices was observed in the PfB9 propeller, which was not found in CyRPA and in most *Plasmodium* B9 proteins (such as PbB9 and PyB9; **Supplementary Fig. 6**). This partially structured region is supported by intrinsic disorder prediction (**Supplementary Fig. 4**), in line with another characteristic of CyRPA, where the loop located on blade 5 likely becomes disordered to accommodate occupancy by a helix of Rh5^43^. The model superimposed well with the CyRPA structure, except for some blade- and strand-connecting loops (RMSD: 3.8 Å; **Fig. 4c**). This similar fold, in addition to the binding activities of CyRPA (targeting Rh5 and Ripr^43^), suggests that the B9 propeller may promote protein-protein interactions.

Since CyRPA is functionally annotated and its binding properties are known, we checked whether the B9 propeller and CyRPA shared a common evolutionary history, which could help to predict the functional sites in the B9 propeller. For this, we generated two datasets consisting of distinct *Plasmodium* B9 (n = 23) or CyRPA (n = 18) sequences (**Supplementary Tables 1** and **2**). Multiple sequence alignments and corresponding phylogenetic trees of these datasets (**Supplementary Files 2** and **3**, and **Supplementary Fig. 7**) were then used concomitantly with their respective tertiary structures to estimate spatially correlated site-specific substitution rates using the GP4Rate tool (**Supplementary Tables 3** and **4**). The six blades were found to be heterogeneously conserved over time for both B9 and CyRPA (Kruskal-Wallis *H* test: B9: *p* = 0.01; CyRPA: *p* = 2.4e-8; **Fig. 4d**). Interestingly, we noticed distinct patterns of evolution between the two proteins: the most conserved blades of B9 propeller (3 and 4) are the less conserved ones in CyRPA (**Fig. 4d**). Because CyRPA interacts with Ripr through its most conserved blade^43^, i.e. 6 (**Fig. 4d**), we logically hypothesize that the blades 3 and 4 of the B9 propeller may target putative partners. Finally, in concordance with different evolutionary histories, we note that the PfB9 propeller and CyRPA display a dissimilar electrostatic surface potential. While almost the entire surface of CyRPA (including the regions mediating interactions with Rh5 and Ripr) is electronegative, some parts of the PfB9 propeller are electropositive (**Fig. 4e**), thus suggesting different binding properties.

### The propeller domain is required for B9 function

We next sought to define the functional importance of the predicted propeller domain, using a structure-guided genetic complementation strategy to evaluate the functionality of truncated B9 proteins (**Fig. 5a**). We assembled various constructs encoding the entire or partially deleted B9, all containing an intact signal peptide and C-terminus sequences to ensure correct secretion and GPI-anchoring of the protein (**Fig. 5b**).

**Figure 5.**
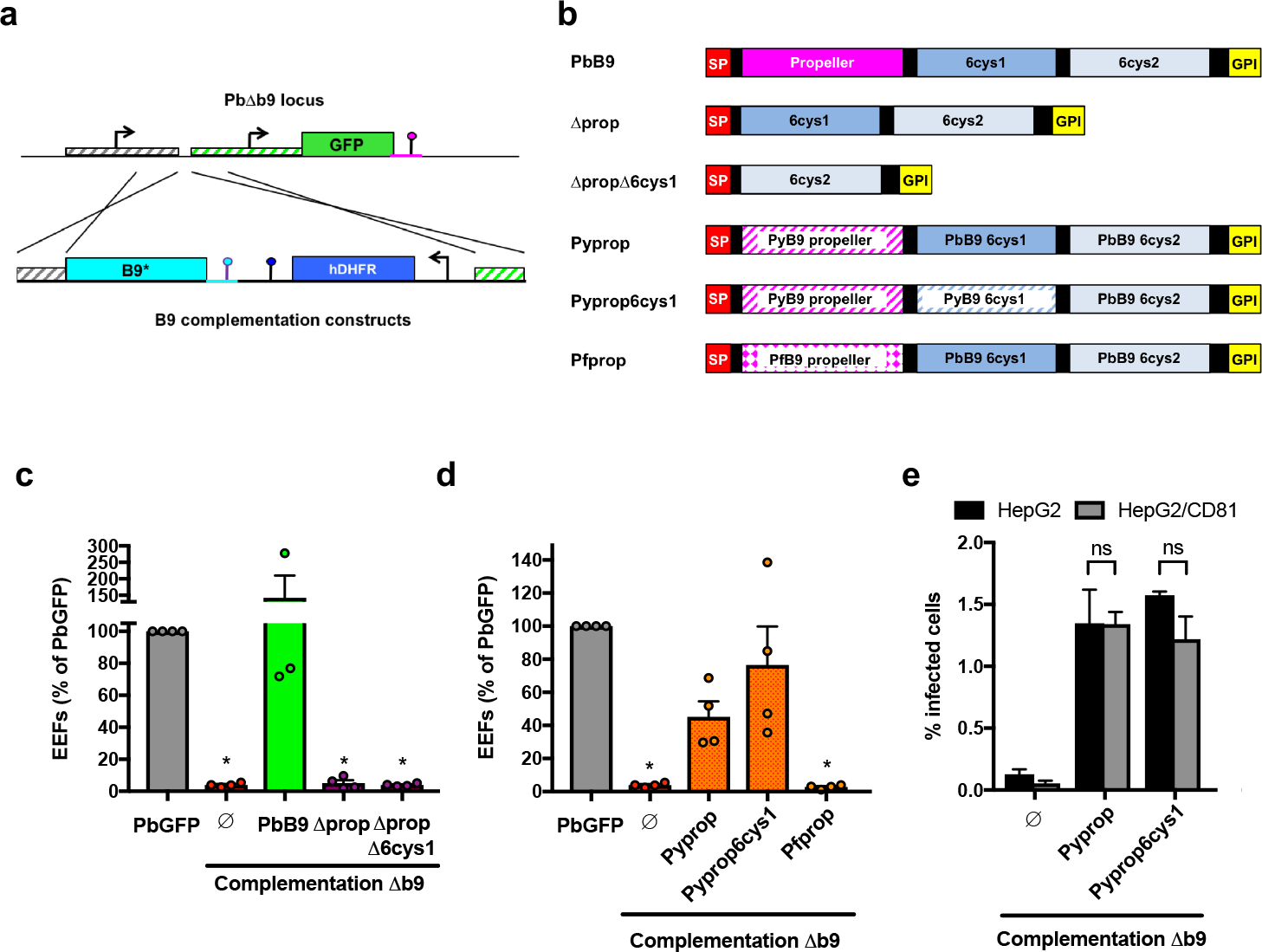
The propeller domain is required for B9 function during sporozoite invasion. **a**, Strategy used to genetically complement PbΔ*b9* with different versions of B9 (indicated as B9*) by double crossover homologous recombination. **b**, Schematic representation of the B9 constructs used for genetic complementation. SP, signal peptide, GPI, glycosylphosphatidylinositol. **c**, Infection rates were determined by quantification of EEFs (GFP-positive cells) 24 h after infection of HepG2 cell cultures with sporozoites of PbGFP, PbΔ*b9* or PbΔ*b9* complemented with PbB9, Δprop or ΔpropΔ6cys1 constructs. Results are expressed as % of control (PbGFP). *p < 0.05 as compared to PbGFP (one-way ANOVA followed by Dunnett’s multiple comparisons test). **d**, Infection rates were determined by quantification of EEFs (GFP-positive cells) 24 h after infection of HepG2 cell cultures with sporozoites of PbGFP, PbΔ*b9* or PbΔ*b9* complemented with Pyprop, Pyprop6cys1 or Pfprop constructs. Results are expressed as % of control (PbGFP). *p < 0.05 as compared to PbGFP (one-way ANOVA followed by Dunnett’s multiple comparisons test). **e**, Infection rates in HepG2 or HepG2/CD81 cells infected with PbΔ*b9* or PbΔ*b9* complemented with PyProp or Pyprop6cys1 constructs were determined 24 hours post-infection. The results show the percentage of invaded (GFP-positive) cells as determined by FACS (mean +/- SD). ns, non-significant (one-way ANOVA followed by Dunnett’s multiple comparisons test).

Constructs were used for transfection of the drug selectable marker-free PbΔ*b9* mutant line. After confirmation of correct integration by genotyping PCR (**Supplementary Fig. 8**), genetically complemented parasites were transmitted to mosquitoes, and sporozoites were tested for infectivity in cell cultures. Complementation of PbΔ*b9* sporozoites with a construct encoding the entire PbB9 fully restored sporozoite infectivity in HepG2 cell cultures (**Fig. 5c**), validating the genetic complementation approach. In contrast, parasites complemented with a truncated B9 lacking the propeller domain, alone or in combination with the first 6-cys domain, were not infectious, phenocopying the parental B9-deficient parasites (**Fig. 5c**). These results show that the propeller domain is required for B9 function during sporozoite entry. Interestingly, chimeric B9 versions where the propeller domain of PbB9 was replaced by the equivalent sequence from PyB9 (Pyprop, Pyprop6cys1; **Fig. 5b**) restored sporozoite infectivity (**Fig. 5d**). In contrast, substitution of the PfB9 propeller domain for the PbB9 propeller (Pfprop; **Fig. 5b**) did not restore infectivity in complemented parasites (**Fig. 5d**). Interestingly, complementation with the PyB9 propeller domain restored infection in both HepG2, which express SR-B1 but not CD81, and HepG2/CD81 cells, which express both receptors^6^, suggesting that the B9 propeller domain does not restrict host receptor usage (**Fig. 5e**).

### The propeller domain of B9 interacts with P36 and P52

Our structural modelling revealed that B9 contains an N-terminus beta-propeller domain structurally similar to CyRPA. In *P. falciparum* merozoites, CyRPA interacts with Rh5 and Ripr to form a complex that is essential for invasion of erythrocytes^42, 43, 45^. While Ripr is conserved among *Plasmodium* species, CyRPA is found in primate but not rodent parasites, and Rh5 is restricted to *P. falciparum* and other *Laverania* species^46^. As Rh5 and Ripr are not expressed by sporozoites^21, 23, 24^, we hypothesized that B9 might be involved in the formation of distinct protein complexes in sporozoites. To test this hypothesis, we first performed co-immunoprecipitation experiments with anti-Flag antibodies, using protein extracts from B9-Flag sporozoites, followed by protein identification by mass spectrometry. However, B9 was the only protein consistently identified in three independent biological replicates by mass spectrometry (data not shown).

We considered that B9 might interact with other sporozoite proteins only at the time of host cell invasion, similarly to CyRPA, which interacts with Rh5 following secretion of merozoite apical organelles^42^. Because sporozoite invasion is a rare event that is difficult to address experimentally, we opted for an alternative strategy based on heterologous expression of sporozoite proteins in mammalian cells, to test for potential interactions between B9 and the 6-cys proteins P36 and P52 as candidate partners, a choice motivated by the shared phenotype of gene deletion mutants. For this purpose, we used a surface display approach to express *P. berghei* proteins at the surface of Hepa1-6 cells after transient transfection^47^. Codon-optimized versions of the propeller domain of PbB9 (amino acids 31-348) or the tandem 6-cys domains of PbP36 (amino acids 67-352) were fused at the N-terminus to the signal peptide of the bee venom melittin (BVM), and at the C-terminus to a V5 epitope tag and the transmembrane domain of glycophorin A, followed by mCherry, C-Myc and 6xHis tags (**Fig. 6a**). As a control, we used an mCherry construct containing all elements except the B9 or P36 sequences. Codon-optimized versions of the tandem 6-cys domains of *P. berghei* P36 and P52 (amino acids 33-302) were expressed either as transmembrane proteins with 3xFlag and GFP tags, or as soluble secreted proteins (sol), with a 3xFlag epitope tag only at the C-terminus (**Fig. 6a**). Interaction between proteins was then tested in co-transfection experiments, by immunoprecipitation followed by western blot. Both P52-GFP (**Fig. 6b**) and P52-sol (**Fig. 6c**) proteins were co-immunoprecipitated with P36-mCherry but not with the control mCherry protein, validating the strategy and confirming the interaction between *P. berghei* P36 and P52 proteins. More importantly, these experiments showed that P36 and P52 co-immunoprecipitated with B9-mCherry, in both transmembrane (**Fig. 6b**) and soluble (**Fig. 6c**) configurations. These results strongly suggest that B9, P36 and P52 form a supramolecular protein complex that, when considering the functional data in this study, appears to mediate productive invasion of hepatocytes by sporozoites.

**Figure 6.**
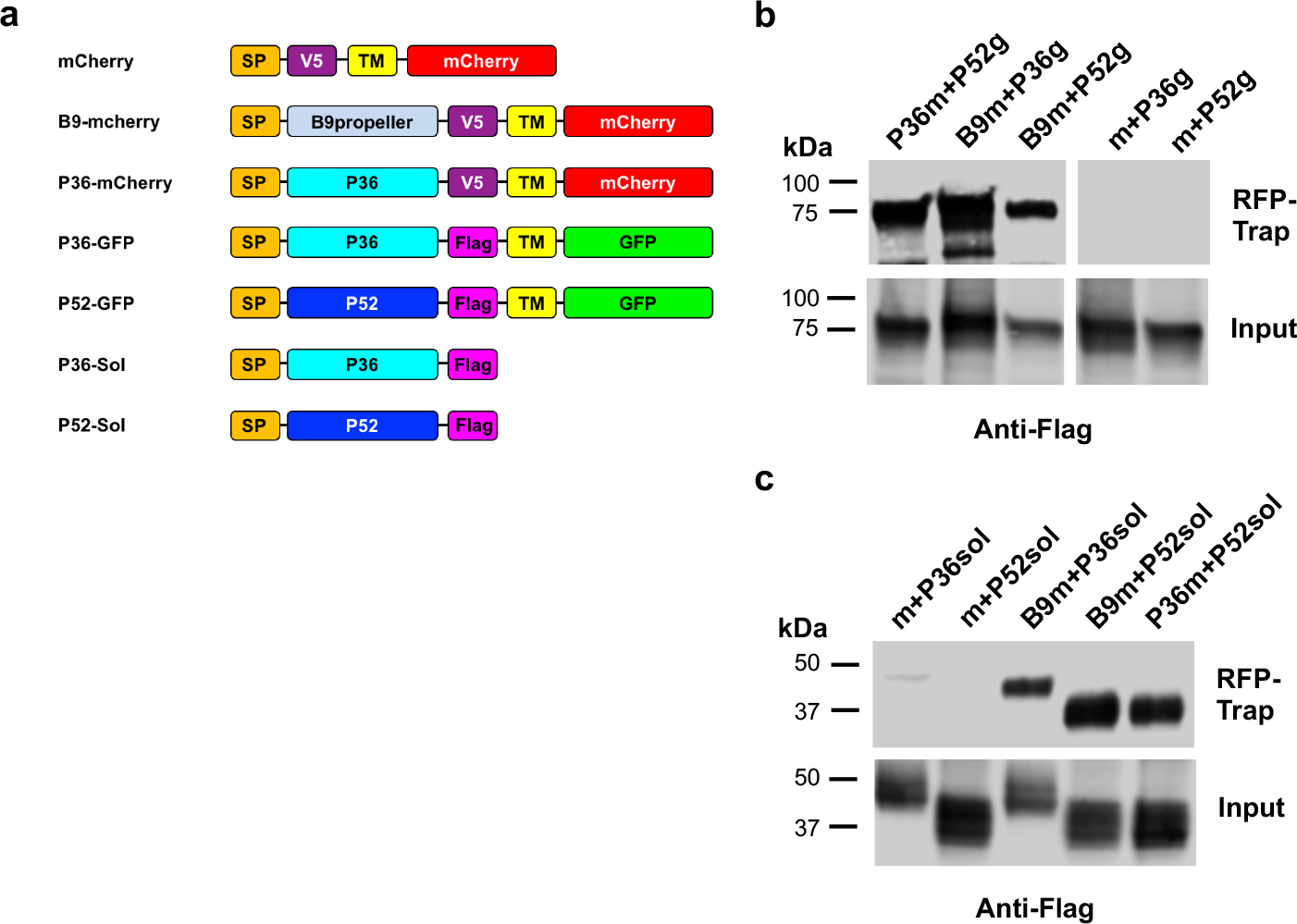
The propeller domain of B9 interacts with P36 and P52. **a**, Schematic representation of the constructs used for heterologous expression in mammalian cells. SP, signal peptide from the bee venom melittin; TM, transmembrane domain and C-terminal portion of mouse Glycophorin A. **b-c**, Hepa1-6 cells were transiently transfected with constructs encoding mCherry (m), B9-mCherry (B9m), or P36-mCherry (P36m) constructs, together with P36-GFP (P36g), P52-GFP (P52g), P36sol or P52sol constructs. Following immunoprecipitation of mCherry-tagged proteins, co-immunoprecipitated proteins (RFP-trap) and total extracts (input) were analysed by western blot using anti-Flag antibodies. The data shown are representative of three independent experiments.

## DISCUSSION

Productive invasion of hepatocytes is a crucial step following transmission of the malaria parasite by a mosquito, however the molecular mechanisms involved remain poorly understood. Until now, only two sporozoite-specific proteins, the 6-cys proteins P36 and P52, have been associated with productive host cell invasion^6, 8^. Here we identify another member of the 6-cys family, B9, as a crucial entry factor. Our data contradict in part those from a previous study, where the authors concluded that B9 is not expressed in sporozoites and is not involved in parasite entry, based mainly on indirect assays^15^. In this study, we demonstrate through genetic tagging that B9 is expressed in *P. berghei* sporozoites, corroborating mass spectrometry data^21–24^. Direct quantification of invasion by flow cytometry established that PbΔ*b9* parasites have an invasion defect. In addition, PbΔ*b9* sporozoites do not discharge their rhoptries upon contact with host cells, similar to PbΔ*p36* sporozoites, indicating that both proteins are acting at an early step during invasion. We further provide evidence that B9 interacts with P36 and P52, suggesting that the three proteins participate in an invasion complex that is required for productive invasion of hepatocytes.

Comparison of profile hidden Markov models between PfB9 and tertiary structure database identified an N-terminus beta propeller domain structurally similar to CyRPA, a cysteine-rich protein expressed in *P. falciparum* merozoites, where it forms a protein complex that is essential for invasion of erythrocytes^42, 43^. Our data suggest that the propeller domain of B9 directly interacts with both P36 and P52. We speculate that blades 3 and 4 of the propeller, which are the most conserved, could be involved in these interactions. Interestingly, the interaction of B9 with P36 and P52 was detected using a heterologous expression system but not by co-IP from sporozoite protein extracts. This observation suggests that B9 may interact with P36 and P52 only after parasite activation, similar to CyRPA, which forms a complex with Rh5 and Ripr only at the time of merozoite invasion in *P. falciparum*^42^. B9 was secreted from sporozoites upon stimulation of microneme exocytosis, as described previously with P36 in *P. yoelii*^11^. B9 shedding could be associated with proteolytic processing, as suggested by the differential migration pattern in western blots. This suggests two possible models, where B9 may bind to P36/P52 either as a membrane-bound or as a free form. Interestingly, although B9 displayed a typical micronemal distribution, it did not colocalize with AMA1. This observation supports the hypothesis that sporozoites contain discrete subsets of micronemes, associated with specific functions^4^.

*P. berghei* and *P. yoelii* sporozoites use different pathways to invade hepatocytes, with the latter being strictly dependent on CD81, like *P. falciparum*^5, 7^. Inter-species complementation experiments have shown that P36 (but not P52) is a key determinant of this differential usage of host receptors^6^. Using a similar approach, we show that the propeller domain of PyB9 can functionally replace the homologous sequence in PbB9, however without altering host receptor usage. This suggests that the B9 propeller does not directly participate in interaction with host receptors. Rather, we hypothesize that B9 may regulate the trafficking and/or binding of P36 to host cells, possibly by concentrating P36-P52 complexes at the surface of the parasite. In contrast, substituting the PfB9 propeller for the *P. berghei* domain abolished protein function, possibly because of impaired interactions with *P. berghei* P36 and/or P52. In this regard, the PfB9 and PbB9 propeller domains show only 48% identity at the amino acid level, versus 90% between PyB9 and PbB9 domains (**Supplementary Fig. 6**). Reciprocally, the essential role of B9 in assembling invasion complexes with P36 and P52 could also explain why *P. falciparum* and *P. vivax* P36 and P52 failed to compensate for the absence of their counterparts in *P. berghei*^6^, as these proteins may not associate with PbB9 to form functional complexes.

Interestingly, an improved version of the neural network-based model AlphaFold^48^ predicts that the C-terminus portion of B9 is organized in three beta sandwiches rather than two (https://alphafold.ebi.ac.uk/). The structures of these domains and their function remain to be experimentally determined. While our data suggest that B9 6-cys-like domains are not required for interaction with P36 and P52, they might regulate the activity of the propeller and/or participate in interactions with host cell surface molecules.

In conclusion, this study reveals that the 6-Cys protein B9 is required for productive host cell invasion by sporozoites. B9 contains a functionally important beta-propeller domain that is likely involved in the formation of a supramolecular protein complex with P36 and P52. Our results suggest that *Plasmodium* sporozoites and merozoites, despite using distinct sets of parasite and host entry factors, may share common structural modules to assemble protein complexes for invasion of host cells. The complex formed by B9, P36 and P52 proteins may represent a potential target for intervention strategies to prevent the initial stages of malaria liver infection.

## METHODS

### Ethics Statement

All mouse work was conducted in strict accordance with the Directive 2010/63/EU of the European Parliament and Council ‘On the protection of animals used for scientific purposes’. Protocols were approved by the Ethical Committee Charles Darwin N 005 (approval #7475-2016110315516522).

### Experimental animals, parasites, and cell lines

*P. berghei* and *P. yoelii* blood stage parasites were propagated in female Swiss mice (6–8 weeks old, from Janvier Labs). We used wild type *P. berghei* (ANKA strain, clone 15cy1) and *P. yoelii* (17XNL strain, clone 1.1), and GFP-expressing PyGFP and PbGFP parasite lines, obtained after integration of a GFP expression cassette at the dispensable *p230p* locus^25^. *Anopheles stephensi* mosquitoes were fed on *P. berghei* or *P. yoelii*-infected mice using standard methods^49^, and kept at 21°C and 24°C, respectively. *P. berghei* and *P. yoelii* sporozoites were collected from the salivary glands of infected mosquitoes 21–28 or 14–18 days post-feeding, respectively. *P. berghei* and *P. yoelii* sporozoite infections were performed in female C57BL/6 or BALB/c mice, respectively (6 weeks old, from Janvier Labs), by intravenous injection in a tail vein. HepG2 (ATCC HB-8065), HepG2/CD81^40^ and Hepa1-6 cells (ATCC CRL-1830) were cultured at 37°C under 5% CO_2_ in DMEM supplemented with 10% fetal calf serum and antibiotics (Life Technologies), as described^7^. HepG2 and HepG2/CD81 were cultured in culture dishes coated with rat tail collagen I (Becton-Dickinson).

### Gene deletion of *p12p, p38* and *b9* in *P. berghei* and *P. yoelii*

Gene deletion mutant parasites were generated using our ‘‘Gene Out Marker Out’’ (GOMO) strategy^25^. For each target gene, a 5’ fragment and a 3’ fragment were amplified by PCR from *P. berghei* (ANKA) or *P. yoelii* (17XNL) WT genomic DNA, using primers listed in **Supplementary Table 5**, and inserted into *Sac*II/*Not*I and *Xho*I/*Kpn*I restriction sites, respectively, of the GOMO-GFP vector^25^, using the In-Fusion HD Cloning Kit (Clontech). The resulting targeting constructs were linearized with *Sac*II and *Kpn*I before transfection. All constructs used in this study were verified by DNA sequencing (Eurofins Genomics). Purified schizonts of *P. berghei* ANKA or *P. yoelii* 17XNL WT parasites were transfected with targeting constructs by electroporation using the AMAXA Nucleofector^TM^ device, as described^50^, and immediately injected intravenously in mice. GFP-expressing parasite mutants were then isolated by flow cytometry after positive and negative selection rounds, as described^25^. Parasite genomic DNA was extracted using the Purelink Genomic DNA Kit (Invitrogen), and analysed by PCR using primer combinations specific for WT, 5’ or 3’ recombined and marker excised loci (listed in **Supplementary Table 6**).

### Genetic tagging of RON4 and B9

Fusion of mCherry at the C-terminus of RON4 was achieved through double crosser homologous recombination. For this purpose, 5’ and 3’ homology fragments, consisting of a 1.2 kb terminal RON4 fragment (immediately upstream of the stop codon) and a 0.6 kb downstream fragment were amplified by PCR using primers listed in **Supplementary Table 7**, and cloned into *Not*I/*Spe*I and *Hind*III/*Kpn*I sites, respectively, of the B3D+mCherry plasmid^51^. The resulting construct was linearized with *Not*I and *Kpn*I before transfection of PbGFP, PbΔ*b9* or PbΔ*p36* purified schizonts. Recombinant parasites were selected with pyrimethamine and cloned by limiting dilution and injection into mice. Integration of the construct was confirmed by PCR on genomic DNA using specific primer combinations listed in **Supplementary Table 7**. Tagging of *P. berghei* B9 with a triple Flag epitope was achieved by double crossover homologous recombination with the endogenous *B9* gene locus. For this purpose, three inserts were amplified by PCR and sequentially inserted in two steps using the In-Fusion HD Cloning Kit (Clontech). In the first step, a 3’ homology 736-bp fragment was cloned into the *Nhe*I site in a plasmid containing a GFP-2A-hDHFR cassette under control of the *P. yoelii* HSP70 promoter. In the second step, a 5’ homology 759-bp fragment from B9 ORF and a 789-bp fragment comprising a triple Flag epitope, a recodonized B9 C-terminus sequence and the 3’ UTR of PyB9 were inserted into *Kpn*I/*Eco*RI sites of the plasmid. Primers used to assemble the B9 tagging construct and the sequence of the synthetic gene are listed in **Supplementary Table 8**. The resulting construct was linearized with *Kpn*I and *Nhe*I before transfection of WT *P. berghei* (ANKA) parasites. Recombinant parasites were selected with pyrimethamine and cloned by limiting dilution and injection into mice. Integration of the construct was confirmed by PCR on genomic DNA using specific primer combinations listed in **Supplementary Table 8**.

### Structure-guided mutagenesis of *P. berghei* B9

Genetic complementation of PbΔ*b9* parasites was achieved by double crossover homologous recombination using a vector containing a hDHFR cassette and a 3’ homology arm corresponding to the 5’ sequence of the HSP70 promoter of the GFP cassette in the PbΔ*b9* line. First, an 840-bp fragment including the coding sequence for PbB9 N-terminus (amino acids 1-29), and a 1096-bp fragment encoding the C-terminus (amino acids 647-852) followed by the 3’ UTR of PbB9 were sequentially inserted into the plasmid, in *Kpn*I/*Eco*RI sites, resulting in the ΔpropΔ6cys1 construct. Cloning of a 1950-bp fragment of PbB9 gene (including the coding sequence for amino acids 30-646) into *Xho*I/*Kpn*I sites of the ΔpropΔ6cys1 plasmid resulted in the PbB9 construct, encoding the full length PbB9 protein. Cloning of a 912-bp fragment of PbB9 gene (including the coding sequence for amino acids 344-646) into *Xho*I/*Kpn*I sites of the ΔpropΔ6cys1 plasmid resulted in the Δprop construct. Cloning of a 1992-bp fragment from PyB9 gene (including the coding sequence for amino acids 30-653 of PyB9) into *Xho*I/*Kpn*I sites of the ΔpropΔ6cys1 plasmid resulted in the PyProp6cys1 construct. Cloning of a 948-bp fragment from PyB9 gene (encoding amino acids 30-342 of PyB9) and a 903-bp fragment from PbB9 gene (encoding amino acids 346-646 of PbB9) into *Xho*I/*Kpn*I sites of the ΔpropΔ6cys1 plasmid resulted in the PyProp construct. Cloning of a 1071-bp fragment from PfB9 gene (encoding amino acids 25-379 of PfB9) and a 903-bp fragment from PbB9 gene (encoding amino acids 346-646 of PbB9) into *Xho*I/*Kpn*I sites of the ΔpropΔ6cys1 plasmid resulted in the PfProp construct. The primers used to assemble the constructs for genetic complementation are listed in **Supplementary Table 9**. The constructs were linearized with *Nhe*I before transfection of PbΔ*b9* purified schizonts. Recombinant parasites were selected with pyrimethamine. Integration of the constructs was confirmed by PCR on genomic DNA using specific primer combinations listed in **Supplementary Table 9**.

### Sporozoite invasion assays

Host cell invasion by GFP-expressing sporozoites was monitored by flow cytometry^52^. Briefly, hepatoma cells (3 × 10^4^ per well in collagen-coated 96-well plates) were incubated with sporozoites (5 × 10^3^ to 1 × 10^4^ per well). For measurement of cell traversal activity, sporozoites were incubated with cells in the presence of 0.5 mg/ml rhodamine-conjugated dextran (Life Technologies). Three hours post-infection, cell cultures were washed, trypsinized and analysed on a Guava EasyCyte 6/2L bench cytometer equipped with 488 nm and 532 nm lasers (Millipore), for detection of GFP-positive cells and dextran-positive cells, respectively. To assess liver stage development, HepG2 or HepG2/CD81 cells were infected with GFP-expressing sporozoites and cultured for 24-36 hours before analysis either by FACS or by fluorescence microscopy, after fixation with 4% PFA and labeling with antibodies specific for UIS4 (Sicgen).

### Fluorescence microscopy

To visualize RON4-mCherry in transgenic parasites, purified schizonts and sporozoites were deposited on poly-L-lysine coated coverslips and fixed with 4% PFA. GFP and mCherry images were captured on a Zeiss Axio Observer.Z1 fluorescence microscope equipped with a Plan-Apochromat 63×/1.40 Oil DIC M27 objective. Images acquired using the Zen 2012 software (Zeiss) were processed with ImageJ or Photoshop CS6 software (Adobe) for adjustment of contrast. To quantify rhoptry discharge, RON4-mCherry expressing PbGFP, PbΔ*b9* or PbΔ*p36* sporozoites were incubated with HepG2 cells for 3 h at 37°C. After extensive washes to remove extracellular parasites, cultures were trypsinized and cells were examined under a fluorescence microscope to assess for mCherry fluorescence in GFP-expressing intracellular sporozoites. At least 50 intracellular parasites in triplicate wells were examined for each parasite line. The percentage of rhoptry discharge was defined as the proportion of intracellular sporozoites without detectable RON4-mCherry signal. For immunofluorescence analysis of B9-Flag parasites, sporozoites collected from infected mosquito salivary glands were deposited on poly-L-lysine coated coverslips, fixed with 4% PFA and permeabilized with 1% Triton X-100. Parasites were labelled with anti-Flag mouse antibodies (M2 clone, Sigma) and AlexaFluor 594-conjugated secondary antibodies (Life Technologies). Nuclei were stained with Hoechst 77742. For double labelling of B9 and AMA1, we used anti-Flag mouse antibodies (M2 clone, Sigma) and anti-AMA1 rat antibodies^53^ (clone 28G2, Bei Resources), followed by atto647N-conjugated anti-mouse and Alexa-594-conjugated anti-rat antibodies. Coverslips were mounted on glass slides with ProLong™ Diamond Antifade Mountant (Life Technologies). STED imaging was carried out with a 93x glycerol-immersion objective (NA 1.3) on a Leica TCS SP8 STEDX microscope equipped with a White Light Laser. AlexaFluor 594 and atto647N-labelled compartments were excited at 590 or 644 nm, respectively, and depleted with a pulsed 775 nm STED laser. Image frames were acquired sequentially frame by frame at a scan speed of 200 lines/s with an optimal pixel size and a line average of 4 to 8. Deconvolution of STED data was performed using the default deconvolution settings in Huygens Professional Deconvolution software v18.10 (Scientific Volume Imaging) that were estimated from the metadata. Brightness and Contrast were adjusted using Fiji^54^.

### Western blot

B9-Flag sporozoites were isolated from the salivary glands of infected mosquitoes and resuspended in 1X PBS. Microneme secretion was stimulated by incubation for 15 min at 37°C in a buffer containing 1% BSA and 1% ethanol, as described^55^. Pellet and supernatant fractions were then isolated from activated and non-activated (control) sporozoites, resuspended in Laemmli buffer and analysed by SDS-PAGE under non-reducing conditions. Western blotting was performed using primary antibodies against the Flag epitope (M2 clone, Sigma) or against GFP (loading control), and secondary antibodies coupled with Alexa Fluor 680. Membranes were then analysed using the InfraRed Odyssey system (Licor).

### Heterologous expression of *Plasmodium* proteins in Hepa1-6 cells

Two vectors for mammalian cell expression were first assembled in a pEF1α-AcGFP1-N1 backbone. The first one (mCherry) encodes a cassette consisting of the signal peptide from bee venom melittin (BVM), a V5 epitope, the transmembrane and C-terminus of mouse Glycophorin A (GYPA), mCherry, Myc and 6xHis tags. In the second one (GFP), the cassette encodes the signal peptide from BVM, a 3xFlag epitope, the transmembrane and C-terminus of mouse GYPA, and GFP. Codon-optimized versions of PbB9 propeller domain (amino acids 31-348), PbP36 (amino acids 67-352) or PbP52 (amino acids 33-302) were inserted in the mCherry and/or GFP plasmids between the signal peptide and the Flag or V5 epitope tag. Two additional constructs for expression of soluble PbP36 and PbP52 were obtained by adding a stop codon immediately after the 3xFlag epitope. The construct cassette sequences are indicated in **Supplementary Table 10**. High concentration plasmid solutions were produced using XL1-Blue Competent Cells (Agilent) and plasmid extraction was performed using Qiagen Plasmid Maxikit (Qiagen) according to the manufacturer’s recommendations. Plasmid transfection was performed in Hepa1-6 cells using the Lipofectamine 2000 reagent (Life Technologies) according to the manufacturer’s specifications. Following plasmid transfection, cells were cultured for 24 h before lysis in a buffer containing 1% NP40. Protein extracts were then subjected to immunoprecipitation using agarose beads coupled with anti-RFP nanobodies (Chromotek). Eluates were collected and analysed by western blot, using anti-Flag antibodies. Membranes were analysed using the InfraRed Odyssey system (Licor).

### B9 immunoprecipitation and mass spectrometry

Freshly dissected B9-Flag sporozoites were lysed on ice for 30 min in a lysis buffer containing 0.5% w/v NP40 and protease inhibitors. After centrifugation (15,000 × g, 15 min, 4°C), supernatants were incubated with protein G-conjugated sepharose for preclearing overnight. Precleared lysates were subjected to B9-Flag immunoprecipitation using Anti-FLAG M2 Affinity Gel (Sigma) for 2h at 4°C, according to the manufacturer’s protocol. PbGFP parasites with untagged B9 were used as a control and treated in the same fashion. After washes, proteins on beads were eluted in 2X Laemmli and denatured (95°C, 5min). After centrifugation, supernatants were collected for further analysis. Samples were subjected to a short SDS-PAGE migration, and gel pieces were processed for protein trypsin digestion by the DigestProMSi robot (Intavis), as described^21^. Peptide samples were analysed on a timsTOF PRO mass spectrometer (Bruker) coupled to the nanoElute HPLC, as described^21^. Mascot generic files were processed with X!Tandem pipeline (version 0.2.36) using the PlasmoDB_PB_39_PbergheiANKA database, as described^21^.

### Structural analyses of B9 propeller

The secondary structure of PfB9 was predicted by hydrophobic cluster analysis^56^ and using PSIPRED 4.0^57^. Conserved domains were searched using InterPro^58^ and HHpred^59^. Glycosylphosphatidylinositol (GPI) anchors were predicted using the NetGPI tool (https://services.healthtech.dtu.dk/service.php?NetGPI)60. Intrinsic disorder prediction was made using the IUPred2A web server (https://iupred2a.elte.hu/)61. The homology model of PfB9 propeller (amino acids 26 to 386) was built with the X-ray structure at 2.4 Å resolution of CyRPA from *P. falciparum* (PDB ID: 5TIH^44^) using the Robetta web server^62^ (default parameters). The model was refined and energy-minimized using respectively GalaxyRefine^63^ and Yasara^64^, then validated using MolProbity^65^ and Prosa II^66^ (**Supplementary Fig. 5**). Structural alignment of PfB9 propeller and CyRPA was performed using the *MatchMaker* function in UCSF Chimera^67^. Protein electrostatic surface potential was calculated using Adaptive Poisson-Boltzmann Solver (APBS^68^), after determining the per-atom charge and radius of the structure with PDB2PQR v.2.1.1^69^. The Poisson-Boltzmann equation was solved at 298 K using a grid-based method, with solute and solvent dielectric constants fixed at 2 and 78.5, respectively. We used a scale of −5 *kT/e* to +5 *kT/e* to map the electrostatic surface potential in a radius of 1.4 Å. All tertiary structures were visualized and drawn using UCSF Chimera^67^.

### Evolutionary analysis of B9 and CyRPA

The amino acid sequence of PfB9 (PlasmoDB code: PF3D7_0317100) and CyRPA (PF3D7_0423800) were queried against the PlasmoDB database^70^ (release 46) and the NCBI non-redundant protein database using blastp searchs (BLOSUM62 scoring matrix). Twenty-three B9 and eighteen CyRPA sequences were retrieved from distinct *Plasmodium* species. Protein sequence alignments were generated using MAFFT version 7 (default parameters^71^). Output alignments were visually inspected and manually edited with BioEdit v7.2.5. Amino acid positions containing gaps in at least 30% of all sequences were removed. Phylogenetic relationships of B9 and CyRPA amino acid sequences were inferred using the maximum-likelihood method implemented in PhyML v3.0^72^, after determining the best-fitting substitution model using the Smart Model Selection (SMS) package^73^. The nearest neighbour interchange approach was chosen for tree improving, and branch supports were estimated using the approximate likelihood ratio aLRT SH-like method^74^. Site-specific substitution rates were estimated by considering their spatial correlation in tertiary structure using the GP4Rate tool^75^. GP4rate requires an amino acid sequence alignment, a phylogenetic tree and a protein tertiary structure to estimate the conservation level during species evolution and the characteristic length scale (in Å) of spatially correlated site-specific substitution rates. For B9, we used the refined tertiary structure predicted by Robetta, while we chose the X-ray structure resolved at 2.4 Å resolution for CyRPA (PDB ID: 5TIH^44^).

### Statistical analysis

Statistical significance of infection data was assessed by Mann-Whitney test or One-way ANOVA followed by Dunnett’s multiple comparisons test. Survival curves were analyzed using the Log rank Mantel-Cox test. All statistical tests were computed with GraphPad Prism 5 (GraphPad Software). *In vitro* experiments were performed with a minimum of three technical replicates per experiment. Statistical analyses for structural modelling were performed using the computing environment R version 3.5.2 (R Foundation for Statistical Computing).

## ACKNOWLEDGEMENTS

We thank Jean-François Franetich, Maurel Tefit and Thierry Houpert for rearing of mosquitoes. The following reagent was obtained through BEI Resources, NIAID, NIH: Monoclonal Anti-Plasmodium Apical Membrane Antigen 1, Clone 28G2 (produced in vitro), MRA-897A, contributed by Alan W. Thomas. This work was funded by grants from the Laboratoire d’Excellence ParaFrap (ANR-11-LABX-0024), the Agence Nationale de la Recherche (ANR-16-CE15-0004) and the Fondation pour la Recherche Médicale (EQU201903007823). The authors acknowledge the Conseil Régional d’Ile-de-France, Sorbonne Université, the National Institute for Health and Medical Research (INSERM) and the Biology, Health and Agronomy Infrastructure (IBiSA) for funding the timsTOF PRO. ML was supported by a ‘DIM 1Health’ doctoral fellowship awarded by the Conseil Régional d’Ile-de-France.

## SUPPLEMENTARY MATERIAL

**Supplementary Figure 1.**
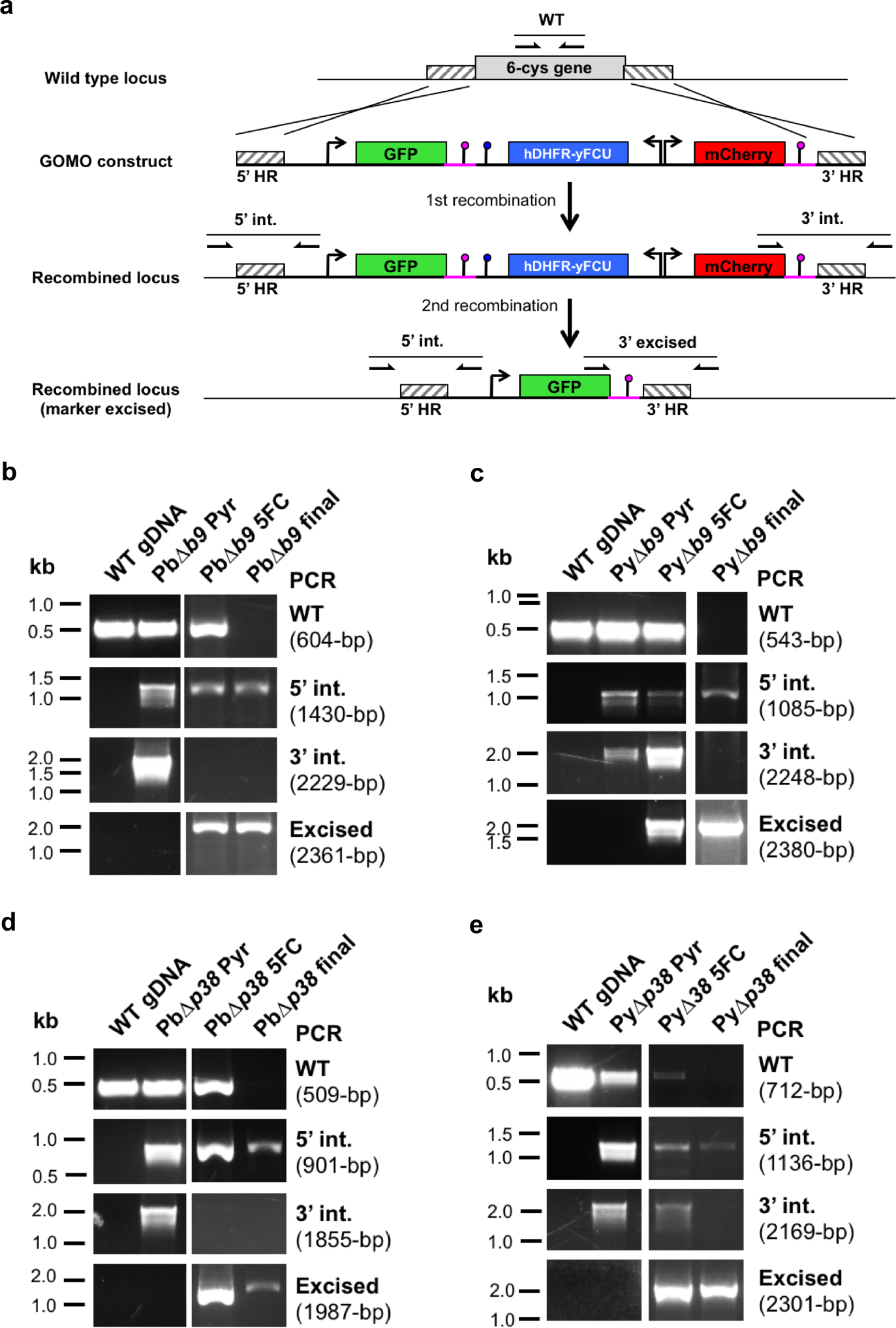
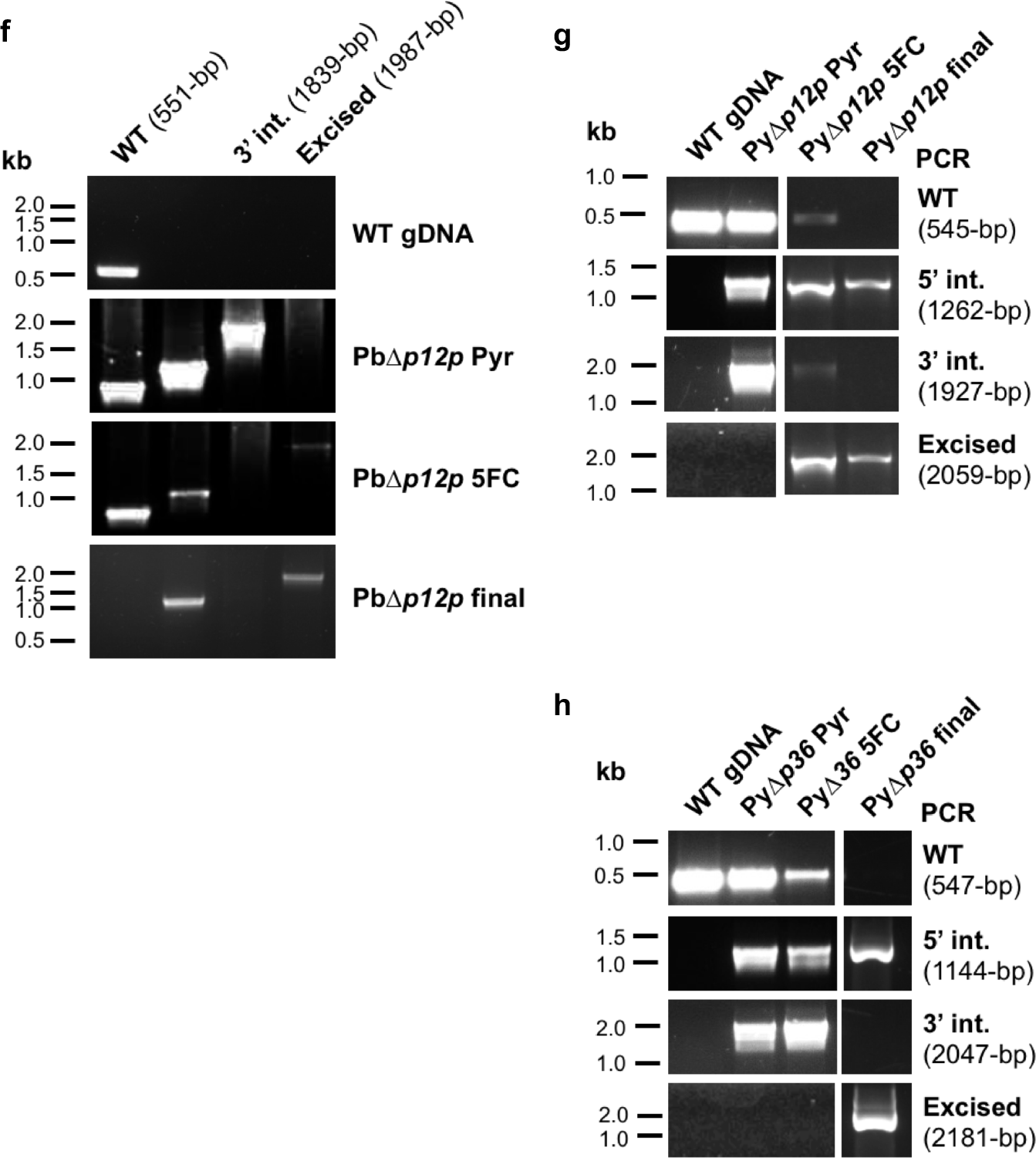
Generation of 6-cys knockout parasite lines in *P. berghei* and *P. yoelii*. **a**, Replacement strategy to delete 6-cys candidate genes. The wild-type locus of 6-cys genes was targeted with a GOMO-GFP replacement plasmid containing a 5’ and a 3’ homologous sequence inserted on each side of a GFP/hDHFR-yFCU/mCherry triple cassette. Upon double crossover recombination, the gene of interest is replaced by the plasmid cassettes. Subsequent recombination between the two identical PbDHFR/TS 3’ UTR sequences (pink lollipops) results in excision of hDHFR-yFCU and mCherry. Genotyping primers and expected PCR fragments are indicated by arrows and lines, respectively. **b–h**, Genotyping of WT and PbΔ*b9* (b), PyΔ*b9* (c), PbΔ*p38* (d), PyΔ*p38* (e), PbΔ*p12p* (f), PbΔ*p12p* (g) and PbΔ*p36* (h) parasites, recovered after positive selection with pyrimethamine (Pyr), negative selection with 5-fluorocytosine (5FC), and parasite sorting by flow cytometry (final). Parasite genomic DNA was analyzed by PCR using primer combinations specific for the unmodified locus (WT), the 5’ integration (5’int.), 3’ integration (3’int.) and 3’ marker excision (excised) events. The absence of amplification with primer combinations specific for the WT locus (WT) and the non-excised integrated construct (3’ integration) confirms that the final populations contain pure knockout drug-selectable marker-free parasites.

**Supplementary Figure 2.**
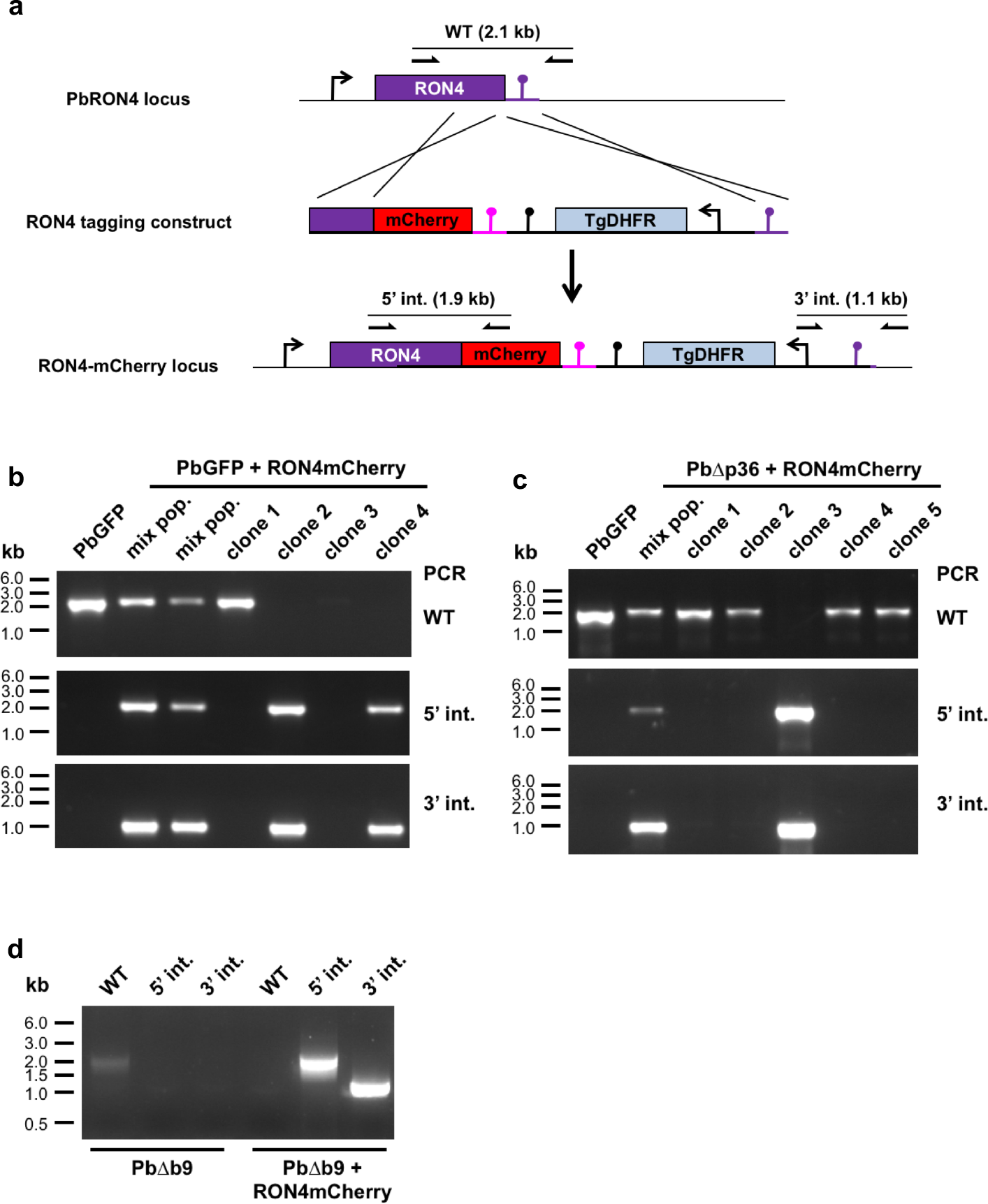
Generation of RON4-mCherry expressing parasites. **a**, Strategy used to tag RON4 with mCherry by double crosser homologous recombination in PbGFP, PbΔ*p36* and PbΔ*b9* parasites. The *P. berghei RON4* locus was targeted with a tagging construct containing a 5’ homology fragment coding the C-terminal part of RON4, fused in frame to the mCherry coding sequence and followed by the 3’ UTR of *P. berghei* DHFR (pink lollipop), a TgDHFR/TS selection cassette, and a 3’ homology fragment corresponding to *RON4* 3’UTR (purple lollipop). Upon a double crossover event, the endogenous *RON4* gene is replaced by a single mCherry-tagged *RON4* copy. Genotyping primers and expected PCR fragments are indicated by arrows and lines, respectively. **b-d**, Correct construct integration was confirmed by analytical PCR using primers specific for the unmodified locus (WT) or for the 5’ and 3’ recombination events (5’ int. and 3’ int., respectively) at the *RON4* locus. The absence of amplification with the WT primer combination confirms the purity of the transgenic population in PbGFP/RON4-mCherry clones 2 and 4 (b), PbΔ*p36*/RON4-mCherry clone 3 (c) and PbΔ*b9*/RON4-mCherry (d).

**Supplementary Figure 3.**
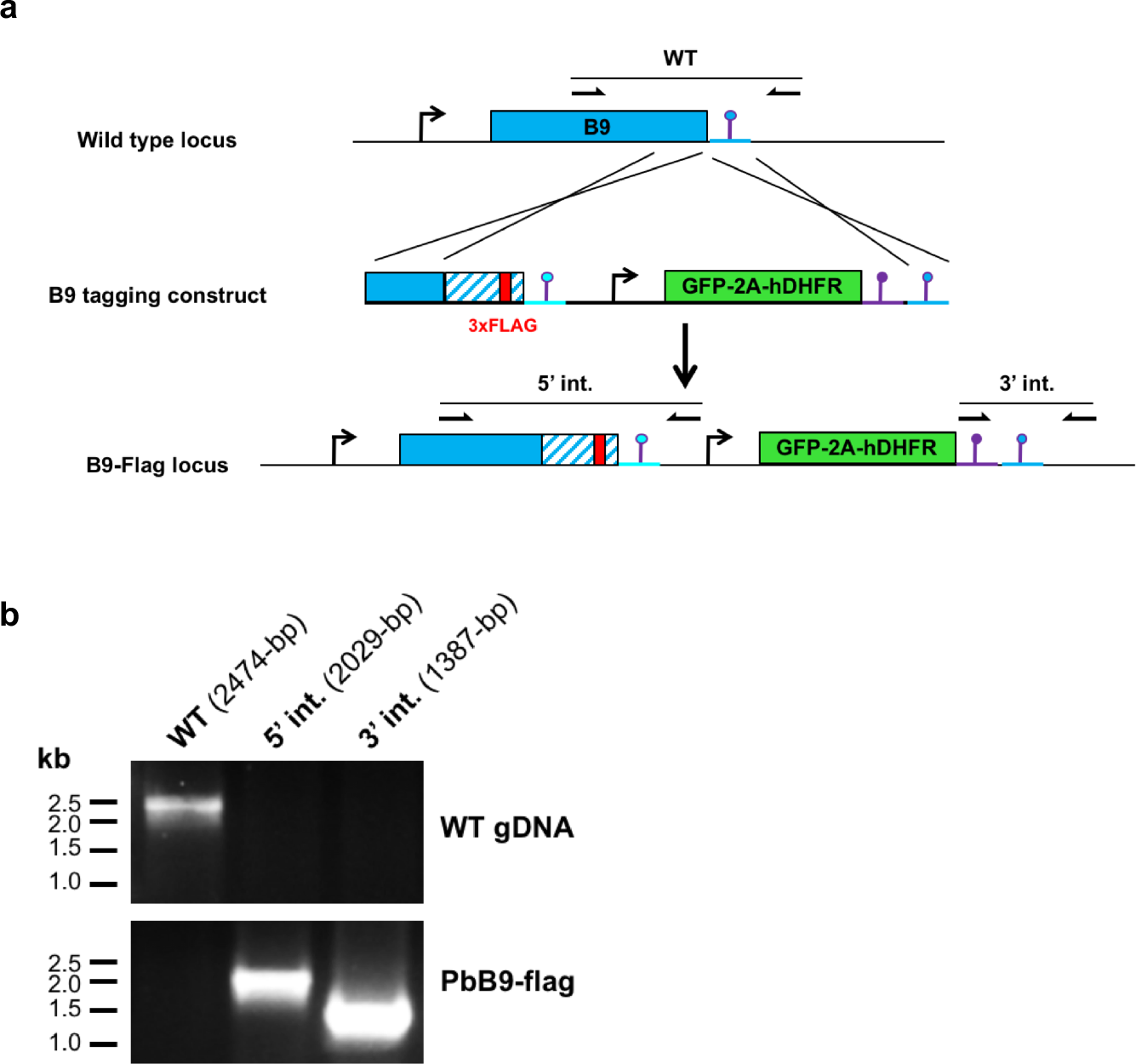
Genetic tagging of B9 in *P. berghei*. **a**, Strategy used to tag B9 with a triple Flag epitope by double crosser homologous recombination in *P. berghei* WT parasites. The *P. berghei B9* locus was targeted with a tagging construct containing a 5’ homology fragment from PbB9 ORF, a recodonized C-terminal sequence of B9 (blue and white striped) with a 3xFlag sequence inserted (red), the 3’ UTR of *PyB9* (cyan lollipop), a GFP-2A-hDHFR cassette, and a 3’ homology fragment corresponding to *PbB9* 3’UTR (blue lollipop). Upon a double crossover event, the endogenous *B9* gene is replaced by a single Flag-tagged *B9* copy. Genotyping primers and expected PCR fragments are indicated by arrows and lines, respectively. **b**, Correct construct integration was confirmed by analytical PCR using primers specific for the unmodified locus (WT) or for the 5’ and 3’ recombination events (5’ int. and 3’ int., respectively) at the *B9* locus. The absence of amplification with the WT primer combination confirms the purity of the transgenic population in PbB9-Flag parasites.

**Supplementary Figure 4.**
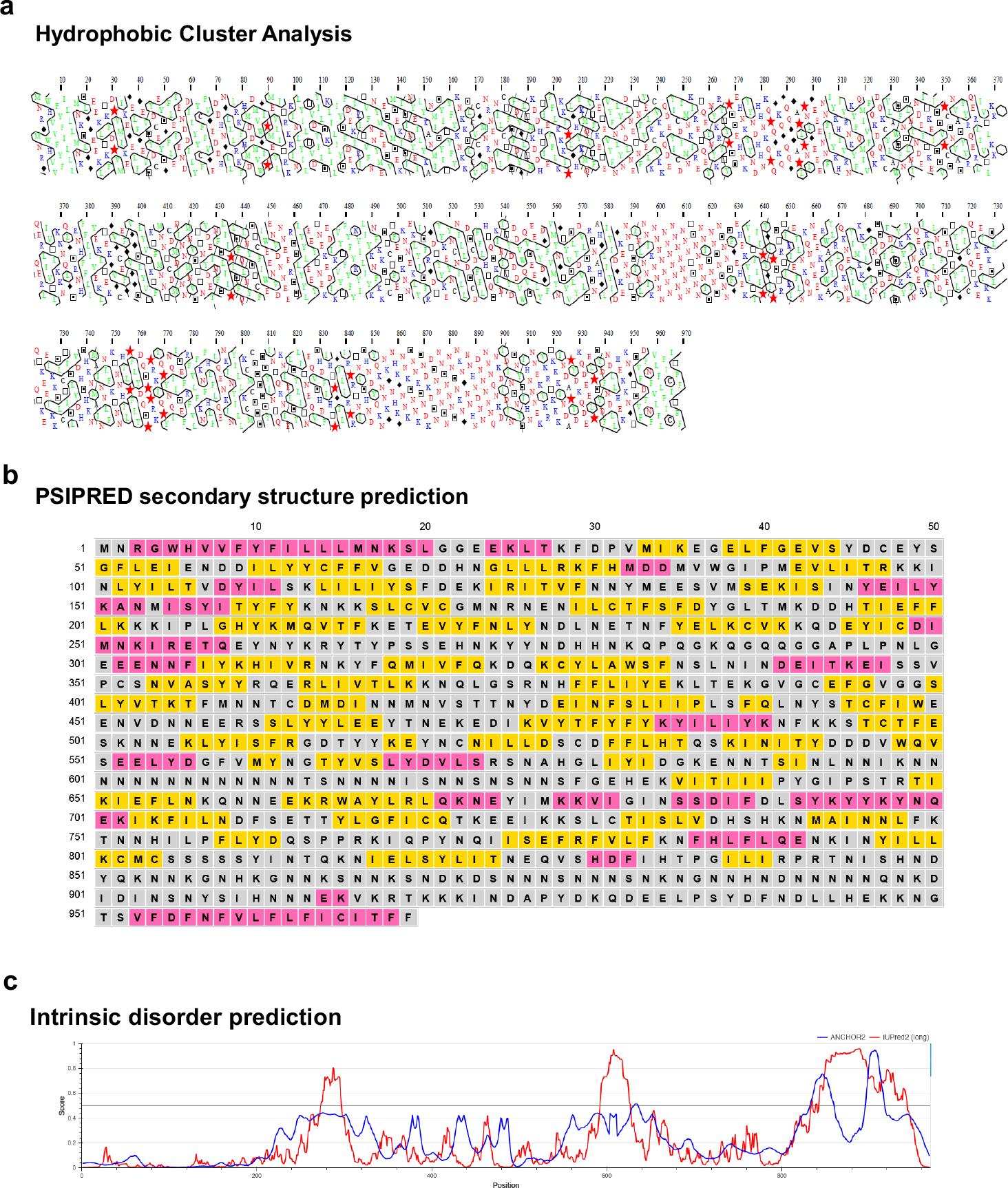
Secondary structure analysis of PfB9. **a**, Hydrophobic cluster analysis of PfB9. Cluster of hydrophobic amino acids are surrounded. Red stars and black diamond correspond to proline and glycine amino acids respectively. **b**, Secondary structure prediction PfB9 using PSIPRED 4.0. Pink, yellow, and grey background colors indicate predicted helix, strand, and coil structures, respectively. **c**. Intrinsic disorder prediction of PfB9 using IUPred2A. Predictions are based on energy estimation for ordered and disordered residues by IUPred2 (red line) and for disordered binding regions by ANCHOR2 (blue line).

**Supplementary Figure 5.**
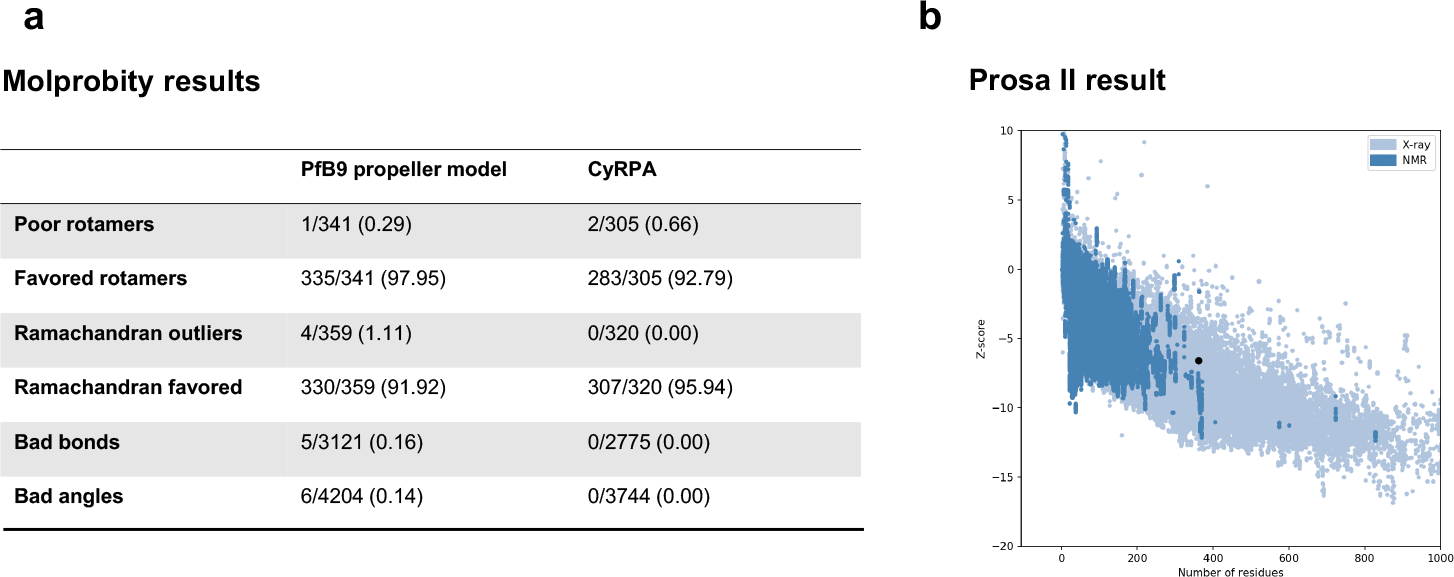
Model structure validation. The structure model of PfB9 propeller (amino acids 26 to 386) was validated using MolProbity (a) and Prosa II (b). For Prosa II, the Z-score of the PfB9 propeller model (indicated as a black point) falls within those of protein structures obtained by X-ray crystallography.

**Supplementary Figure 6.**
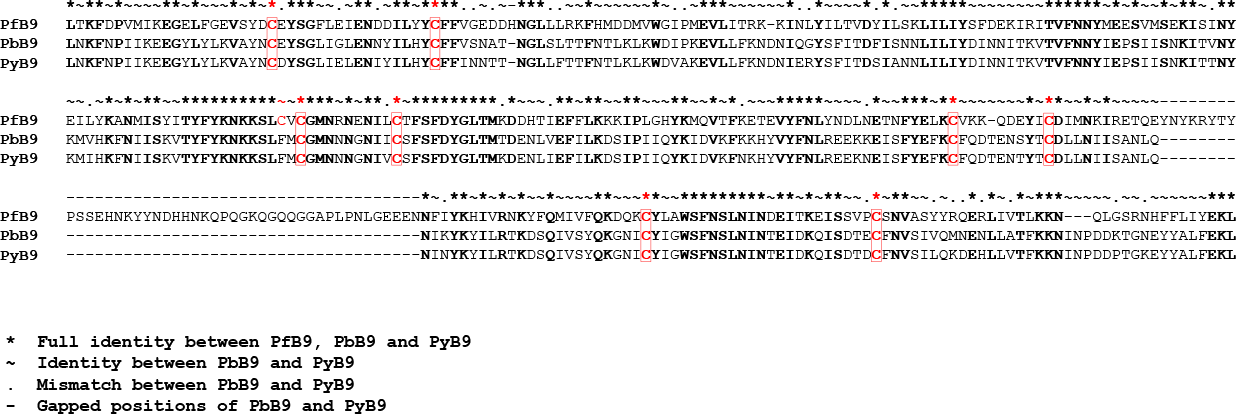
Protein sequence alignment of the B9 propeller from *P. falciparum*, *P. berghei* and *P. yoelii*. Conserved residues are in bold, cysteines in red.

**Supplementary Figure 7.**
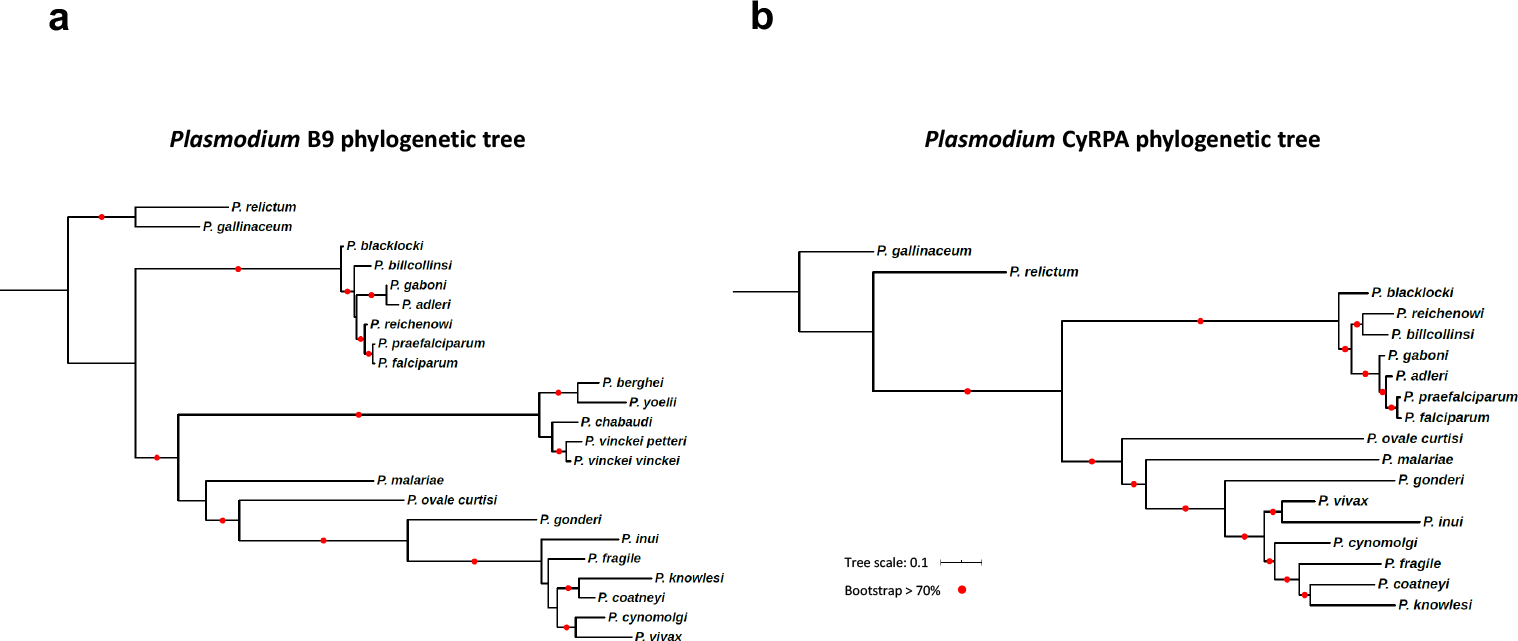
Phylogenetic trees of B9 and CyRPA. Multiple sequence alignments and corresponding phylogenetic trees were established using two datasets consisting of distinct *Plasmodium* B9 (n = 23) or CyRPA (n = 18) sequences. Phylogenetic trees were inferred by maximum likelihood using PhyML, and branch supports were estimated using the approximate likelihood ratio aLRT SH-like method.

**Supplementary Figure 8.**
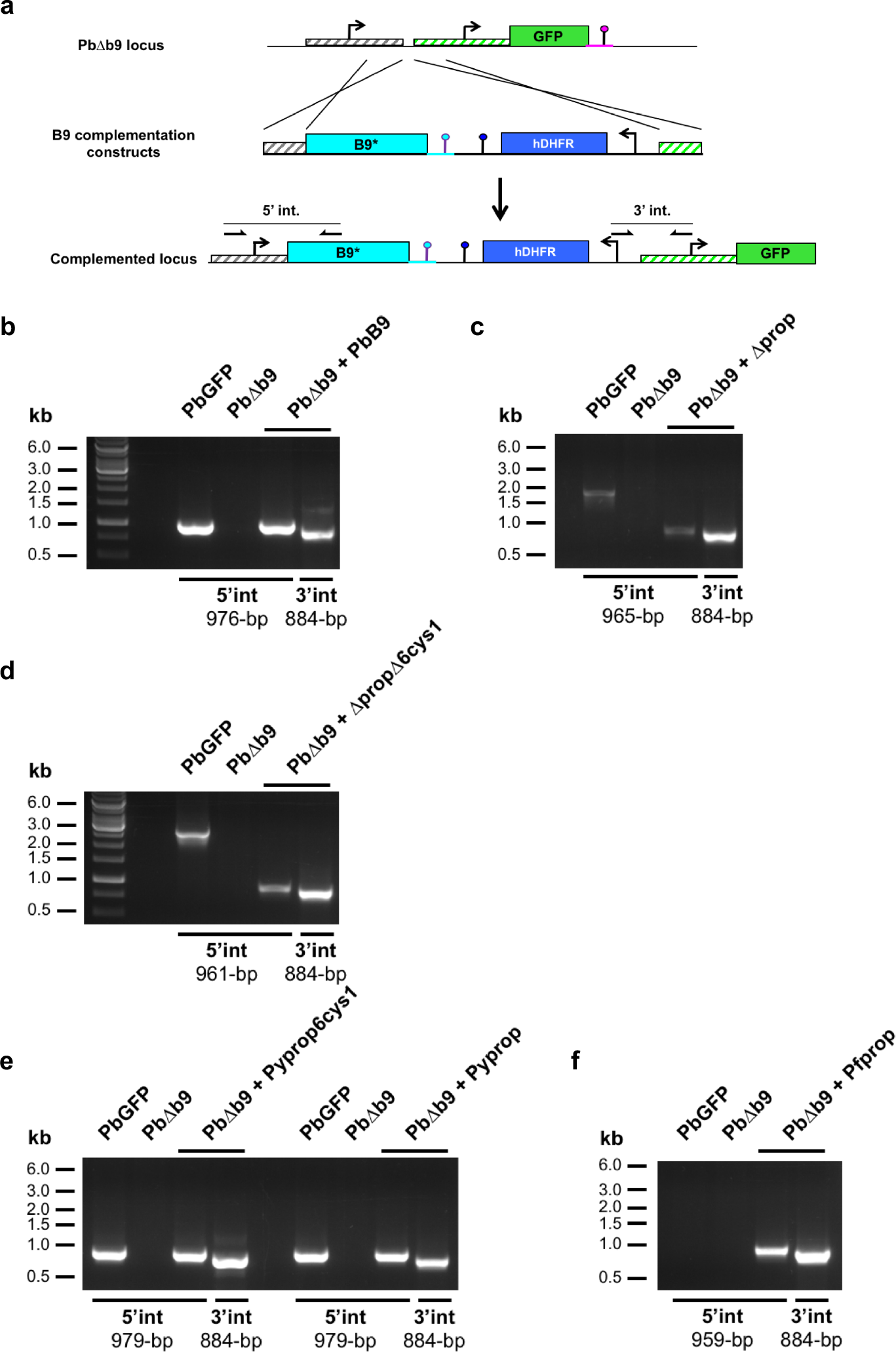
Genetic complementation of PbΔb9 parasites. **a**, Strategy used to genetically complement PbΔ*b9* with different versions of B9 (indicated as B9*) by double crossover homologous recombination. Genotyping primers and expected PCR fragments are indicated by arrows and lines, respectively. **b-f**, Correct construct integration was confirmed by analytical PCR using primers specific for the 5’ and 3’ recombination events (5’ int. and 3’ int., respectively) and genomic DNA from PbGFP, PbΔ*b9*, and PbΔ*b9* complemented with the PbB9 (b), Δprop (c), ΔpropΔ6cys (d), Pyprop (e), Pyprop6cys1 (e) or Pfprop (f) constructs.

